# Pyruvate dehydrogenase kinase 1 controls triacylglycerol hydrolysis in cardiomyocytes

**DOI:** 10.1101/2024.10.14.618123

**Authors:** Michael G. Atser, Chelsea D. Wenyonu, Elyn M. Rowe, Connie L. K. Leung, Haoning Howard Cen, Eric D. Queathem, Leo T. Liu, Renata Moravcova, Jason Rogalski, David Perrin, Peter Crawford, Leonard J. Foster, Armando Alcazar, James D. Johnson

## Abstract

Pyruvate dehydrogenase kinase (PDK) 1 is one of four isozymes that inhibit the oxidative decarboxylation of pyruvate to acetyl-CoA via pyruvate dehydrogenase. PDK activity is elevated in fasting or starvation conditions to conserve carbohydrate reserves. PDK has also been shown to increase mitochondrial fatty acid utilization. In cardiomyocytes, metabolic flexibility is crucial for the fulfillment of high energy requirements. The PDK1 isoform is abundant in cardiomyocytes, but its specific contribution to cardiomyocyte metabolism is unclear. Here we show that PDK1 regulates cardiomyocyte fuel preference by mediating triacylglycerol turnover in differentiated H9c2 myoblasts using lentiviral shRNA to knockdown *Pdk1*. Somewhat surprisingly, PDK1 loss did not affect overall PDH activity, basal glycolysis, or glucose oxidation revealed by oxygen consumption rate experiments and ^13^C_6_ glucose labelling. On the other hand, we observed decreased triacylglycerol turnover in H9c2 cells with PDK1 knockdown, which was accompanied by decreased mitochondrial fatty acid utilization following nutrient deprivation. ^13^C_16_ palmitate tracing of uniformly labelled acyl chains revealed minimal acyl chain shuffling within triacylglycerol, indicating that the triacylglycerol hydrolysis, and not re-esterification, was dysfunctional in PDK1 suppressed cells. Importantly, PDK1 loss did not significantly impact the cellular lipidome or triacylglycerol accumulation following palmitic acid treatment, suggesting that effects of PDK1 on lipid metabolism were specific to the nutrient-deprived state. We validated that PDK1 loss decreased triacylglycerol turnover in *Pdk1* knockout mice. Together, these findings implicate a novel role for PDK1 in lipid metabolism in cardiomyocytes, independent of its canonical roles in glucose metabolism.

## Introduction

Cardiomyocytes must continually produce ATP to keep up with high energy demands because they possess little energy reserves (1). Healthy adult cardiomyocytes meet their high energy requirement by oxidizing various fuel sources such as fatty acids (40-60% of ATP), carbohydrates (glucose and lactate; 20-40% of ATP), amino acids, and ketones (2–4). There is a reciprocal relationship between glucose and fatty acid utilization (2, 5–8). Acetyl-CoA generated from fatty acid β-oxidation can activate pyruvate dehydrogenase kinase (PDK), an inhibitor of the pyruvate dehydrogenase complex (PDH), thereby inhibiting the entry of glucose carbon units into acetyl-CoA. On the other hand, citrate can be converted to acetyl-CoA by citrate lyase in the cytosol, which is then converted to malonyl-CoA to inhibit carnitine palmitoyl transferase I and prevent the rate-limiting mitochondrial import of fatty acids (3, 9–11). Thus, acetyl-CoA is essential for regulating the utilization of glucose and fatty acids, with the PDK/PDH axis serving as a crucial control point in this process (12–14).

PDH is a multiunit complex enzyme that oxidatively decarboxylates pyruvate to acetyl-CoA and CO_2_ while coupling the reduction of NAD^+^ to NADH (15, 16). Since no known pathway exists for converting acetyl-CoA to glucose, fuel source switching to fatty acids or other nutrients is crucial in conserving glucose in the fasted state (13). PDH is regulated via a phosphorylation/dephosphorylation cycle controlled by PDK and pyruvate dehydrogenase phosphatase (PDP) respectively (16).

There are four PDK isozymes (PDK1, PDK2, PDK3, and PDK4) that differ in tissue abundance, regulation and reaction kinetics (13, 17). All PDK isozymes phosphorylate PDH at serine 293 and serine 300. However, PDK1 is the only isoform to phosphorylate PDH on serine 232 (18, 19). Pyruvate allosterically inhibits PDK activity while increasing NADH/NAD^+^ (13). Acetyl-CoA/CoA ratios activate PDK activity (13). PDK1 shows low sensitivity to pyruvate inactivation but is readily responsive to NADH/NAD^+^ and acetyl-CoA/CoA activation (20). At the transcriptional level, *Pdk2* and *Pdk4* mRNA levels are upregulated in response to glucose deprivation with fatty acid supplementation and downregulated on the addition of insulin (15). In contrast, *Pdk1* is resistant to glucose and insulin levels (21–23). During starvation, overall cardiac PDK activity is elevated to inhibit PDH and conserve carbohydrate reserves (16). Though it is abundant in the heart, the specific contribution of PDK1 to this process remains ambiguous.

In humans, rare variants in *PDK1* are associated with multiple metabolic traits, including body fat percentage, heart failure, type 2 diabetes, non-alcoholic fatty liver disease, and circulating pyruvate levels (www.hugeamp.org). Transcriptomics analysis reveal that *PDK1* expression is upregulated in hearts of patients with cardiomyopathies such as arrhythmogenic right ventricular cardiomyopathy and dilated cardiomyopathy (https://ngdc.cncb.ac.cn/cvd/gene/CVDG012320) (24). Although proteomics studies revealed no changes in PDK1 protein abundance (25–28), a recent phosphoproteomics study of diabetic and ischemic cardiomyopathic patients revealed decreased phosphorylation of the PDK1-specific serine 232 site on PDH (29), further implicating PDK1 in cardiac disease. In mice, we and others identified the *Pdk1* gene as a positional and functional candidate in genome-wide association studies of diet-induced obesity (30, 31). Global *Pdk1* knockout mice fed a high fat diet did not show any changes to overall adiposity but exhibited dysregulated cardiac lipid homeostasis in the fasted state (31). These findings implicated a role for PDK1 in cardiomyocyte lipid metabolism, but the specific actions on glucose and fatty acid metabolism were unknown.

In this study, we characterize PDK1’s action on cardiomyocyte glucose and fatty acid metabolism. We found that PDK1 mediates triacylglycerol turnover without altering glucose metabolism in nutrient-deprived cells previously fed excess palmitic acid.

## Results

### Validation of Pdk1 knockdown H9c2 cells

To determine the specific role of the *Pdk1* gene in cardiomyocytes, PDK1 protein abundance was stably suppressed by shRNA lentiviral transduction in H9c2 cells, a surrogate cardiomyocyte cell line.

Cells were transduced with four shRNA targeting different regions of the *Pdk1* mRNA and a non-targeting scrambled shRNA was used as a control. The shRNA packaging plasmids contained a puromycin resistance gene to confer puromycin resistance unto cells with stable genomic integration. Following puromycin treatment, positively selected cells were analyzed for PDK1 suppression. Western blot analysis identified an shRNA with a marked reduction in PDK1 protein abundance (Fig S1A). Scrambled-control (Control) and shRNA transduced (PDK1 knockdown) cells were then differentiated to a cardiomyocyte phenotype in 1% FBS and 1 μM of *trans*-retinoic acid for 9 days (32, 33). Both cell lines showed increased expression of a cardiomyocyte maturation marker, *Tnnt*, validating the differentiations (Fig S1B). To assess the effect of the transduced shRNA on the other PDK isoforms, we measured the protein abundance in differentiated control and PDK1 knockdown (KD) cells using LC/MS-MS proteomics. As expected, we observed a significant decrease in PDK1 in KD cells (Fig 1A). We observed an increase in PDK3 in KD cells, indicative of some compensation (Fig 1A). We were unable to reliably detect *Pdk4* mRNA or protein abundance by qPCR and mass spectrometry, suggesting its levels are low in these cells. Together, our data validate our PDK1 knockdown model in differentiated H9c2 cells.

**Figure 1:**
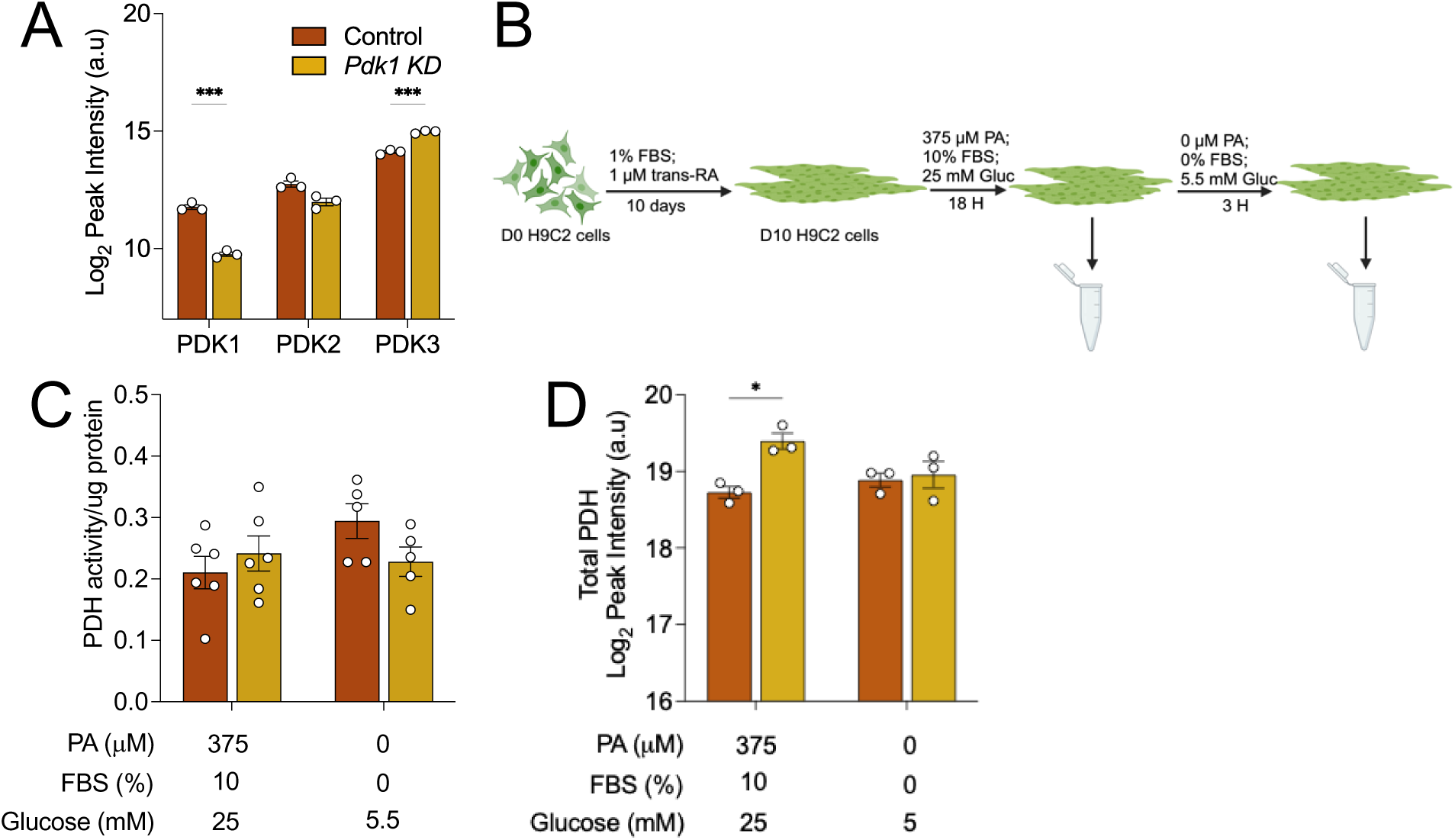
Effect of PDK1 suppression on PDH activity. **(A)** PDK protein abundance measured by LC-MS/MS based proteomics respectively in control and *Pdk1* knockdown (KD) differentiated H9c2 cardiomyocytes. **(B)** Schematic of experimental design showing differentiation of shRNA transduced H9c2 myoblasts for 10 days in 1% fetal bovine serum and 1 μM trans-retinoic acid for 10 days followed by pre-treatment with 375 μM palmitic acid in complete growth medium for 18 h and subsequent nutrient deprivation in low glucose, serum-free media. **(C)** Total PDH activity, **(D)** Total PDH abundance were assessed in control and Pdk1 knockdown cells following palmitic acid treatment and nutrient deprivation. Data are shown as mean ± SEM and were analyzed by either a Two-way ANOVA followed by a post-hoc Tukey multiple comparison test (C, D) or multiple t-test with Bonferroni-Dunn correction (A). For (C), ANOVA showed (F and p values) of (3.1 and 0.0906), (1.6 and 0.2118), (0.4 and 0.526) for interaction, row (nutrient availability) and column (cell lines) factors respectively. For (D), ANOVA showed (F and p) values of (6.523 and 0.0340), (1.416 and 0.2682), (9.783 and 0.0141) for interaction, row (nutrient availability) and column (cell lines) factors respectively. ^∗^ p< 0.05; ^∗∗^ p< 0.01; ^∗∗∗^ p< 0.001; ∗∗∗∗ p<0.0001.

### PDK1 suppression does not alter PDH Activity in differentiated H9c2 cells

PDKs phosphorylate PDH to inhibit its activity and regulate the entry of glucose’ carbon units into the citric acid cycle. In the fed state, glucose is the primary carbon source for mitochondrial oxidation in cardiomyocytes (34). However, during high fatty acid or fasted conditions, fatty acids are the primary carbon source and glucose utilization is inhibited by PDKs (34). To determine if PDK1 specifically is involved this regulation, differentiated control and KD cells were treated with 375 μM of palmitic acid complexed to BSA (6:1 molar ratio) in complete medium (25 mM glucose, 10% FBS) for 18 h to stimulate triacylglycerol accumulation, then nutrient deprived in serum-free, low glucose medium (5.5 mM glucose, 0% FBS) for 3 h (Fig 1B). We observed that total PDH activity was not significantly different between control and KD cells in both the palmitic acid-fed and the nutrient deprived conditions (Fig 1C). Surprisingly, mass spectrometry analysis of PDH protein abundance revealed upregulation in KD cells in the palmitic acid-fed condition with no difference in levels between control and KD in the nutrient deprived condition (Fig 1D). This finding prompted us to look again at the abundance of the other PDK isoforms in these conditions, to determine if a change in abundance may provide insights into these findings. Mass spectrometry based proteomic analysis showed the maintained suppression of PDK1 in KD cells in both conditions, as expected (Fig S1C). Although PDK2 was unchanged in KD cells in both conditions, PDK3 remained elevated in KD cells in only the palmitic acid-fed condition, returning to control levels in the nutrient deprived condition (Fig S1C). These findings suggest that elevation of PDK3 may counteract the elevated PDH protein abundance to maintain PDH activity in the palmitic acid-fed condition.

### PDK1 suppression does not alter glucose utilization in differentiated H9c2 cells

Because PDK1 suppression did not alter total PDH activity in these cells, we next asked if glucose metabolism was impacted in a manner independent of total PDH activity. To assess glucose consumption, we measured the glucose in media following palmitic acid treatment and nutrient deprivation respectively. Predictably, the glucose consumed was higher in the 18 h palmitic acid treated condition than the 3 h nutrient deprived condition across both cell lines presumably due to the longer incubation time in the former (Fig 2A). However, there was no difference between control and KD cells in both conditions (Fig 2A). We also didn’t observe any significant difference in intracellular glucose levels between control and KD in both palmitic acid treated and nutrient deprived conditions (Fig 2B). Interestingly, the intracellular glucose pool was observed to decrease in both cell lines in the nutrient deprived condition (Fig 2B). Despite intracellular glucose being unchanged in the palmitic acid treatment, we observed increased glycogen levels in KD cells with no significant difference between control and KD in the nutrient deprived condition (Fig 2C). We then investigated how elevated glycogen levels in PDK1 suppressed cells might impact glucose metabolism following palmitic acid treatment. We found no difference in basal glycolytic rate but a significant elevation in compensatory glycolytic rate following the inhibition of mitochondrial oxidation (Fig 2D-F). Furthermore, we found no change in the dependency on glucose oxidation but noted an elevation in glucose oxidation capacity when alternative pathways such as fatty acid and amino acid oxidation were inhibited (Fig 2G-J). The rate of ATP production from either glycolysis or mitochondria was also not significantly different (Fig 2K-M).

**Figure 2:**
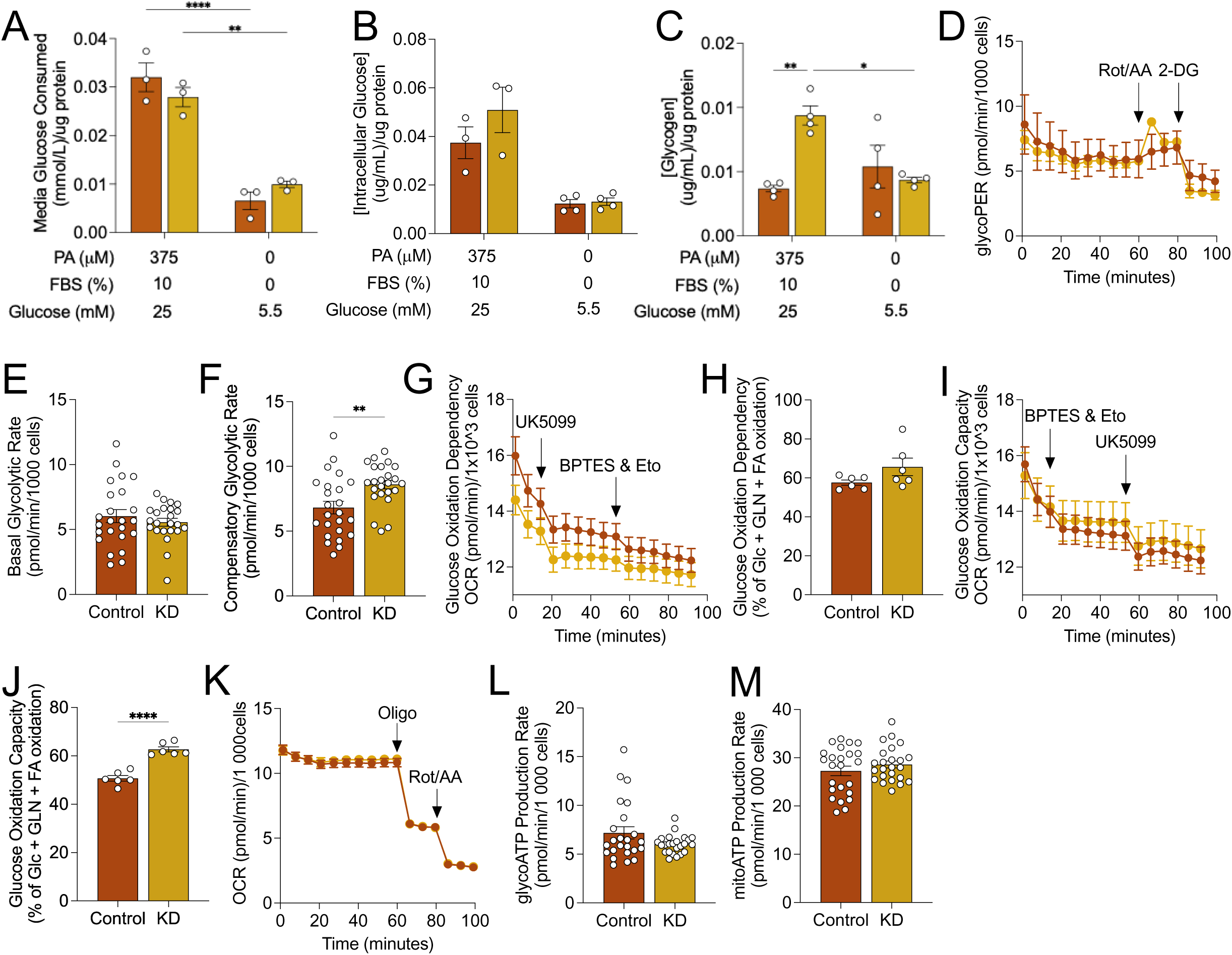
Effect of PDK1 suppression on glucose metabolism. **(A)** Consumed glucose, **(B)** intracellular glucose, **(C)** glycogen abundance was measured in differentiated control and KD cells following 375 μM palmitic acid treatment and nutrient deprivation for 3 h. Control and KD cells were treated with 375 μM palmitic acid treatment for 18 h and **(D)** glycolytic proton efflux rate, **(E)** basal glycolytic rate and **(F)** compensatory glycolytic rate **(G&H)** glucose oxidation dependency, **(I&J)** glucose oxidation capacity and **(K-M)** ATP production rate were measured from independent cell cultures spanning 3 experimental replicates. Data are shown as mean ± SEM and were analyzed by either a Two-way ANOVA followed by a post-hoc Tukey multiple comparison test (A - C) or Two-tailed t-test. For (A), ANOVA showed (F and p values) of (3.375 and 0.1035), (113.8 and <0.0001), (0.02796 and 0.8714) for interaction, row (nutrient availability) and column (cell lines) factors respectively. For (B), ANOVA showed (F and p values) of (1.617 and 0.2323), (40.11 and <0.0001), (2.083 and 0.1796) for interaction, row (nutrient availability) and column (cell lines) factors respectively. For (C), ANOVA showed (F and p values) of (13.28 and 0.0034), (3.203 and 0.0987), (6.358 and 0.0268) for interaction, row (nutrient availability) and column (cell lines) factors respectively. ^∗^ p< 0.05; ^∗∗^ p< 0.01; ^∗∗∗^ p< 0.001; ∗∗∗∗ p<0.0001.

These findings suggest that although basal glucose metabolism may remain unchanged, PDK1 knockdown cells may possess a greater glucose metabolic reserve. We are unsure of the mechanism underlying the link between PDK1 suppression and elevated glycogen in the nutrient-rich condition (Fig 2C). The depletion of this excess glycogen in the nutrient-deprived condition may suggest that the increased glucose oxidation capacity in PDK1 suppressed cells during nutrient deprivation may be due to elevated glycogen stores in the nutrient rich condition.

### PDK1 suppression increases pyruvate flux through malate/oxaloacetate axis

Nutrient oxidation dependency can be influenced by a variety of factors such as the rate of oxidation and the delivery of energy substrates to the mitochondria. Since PDKs regulate the committal step of glucose oxidation, we were curious how the rate of glucose oxidation may be impacted in PDK1 suppressed cells. In addition, *in vitro* measurements of enzymatic activity from cell lysates do not sometimes reflect the biological activity in cells (35). Therefore, we fed control and KD cells with 150 μM palmitic acid for 18 h in complete medium and then nutrient deprived with 5.5 mM of unlabelled glucose for 2 h followed by incubation in 5.5 mM of uniformly labeled [U-^13^C_6_] glucose for 1 h, which is metabolized through the glycolytic pathway (Fig 3A). The relative fractional enrichments of each metabolite are shown in supplementary Figure S2A-I. We found no significant differences in the pool sizes of unlabeled glycolytic or citric acid cycle (TCA) intermediates following nutrient deprivation (Fig S2J). We measured the fractional enrichment of glycolytic intermediates to validate that glycolysis was similar in both cell lines. We observed no significant difference in the fractional enrichment of hexose 6-phosphate with 6 labeled carbons (M+6) and phosphoenolpyruvate with 3 labeled carbons (M+3) (Fig 3B&C). Although, we were unable to detect pyruvate in our spectra due to high noise levels exceeding the limits of detection, we measured the fractional enrichment of M+3 alanine as a proxy for [U-^13^C_3_] pyruvate since it is derived directly from the transamination of pyruvate (36). We observed no significant difference in M+3 alanine enrichment across cell lines (Fig 3D). Taken together, these results validate our prior finding that there is no significant difference in glycolysis between control and KD cells.

**Figure 3:**
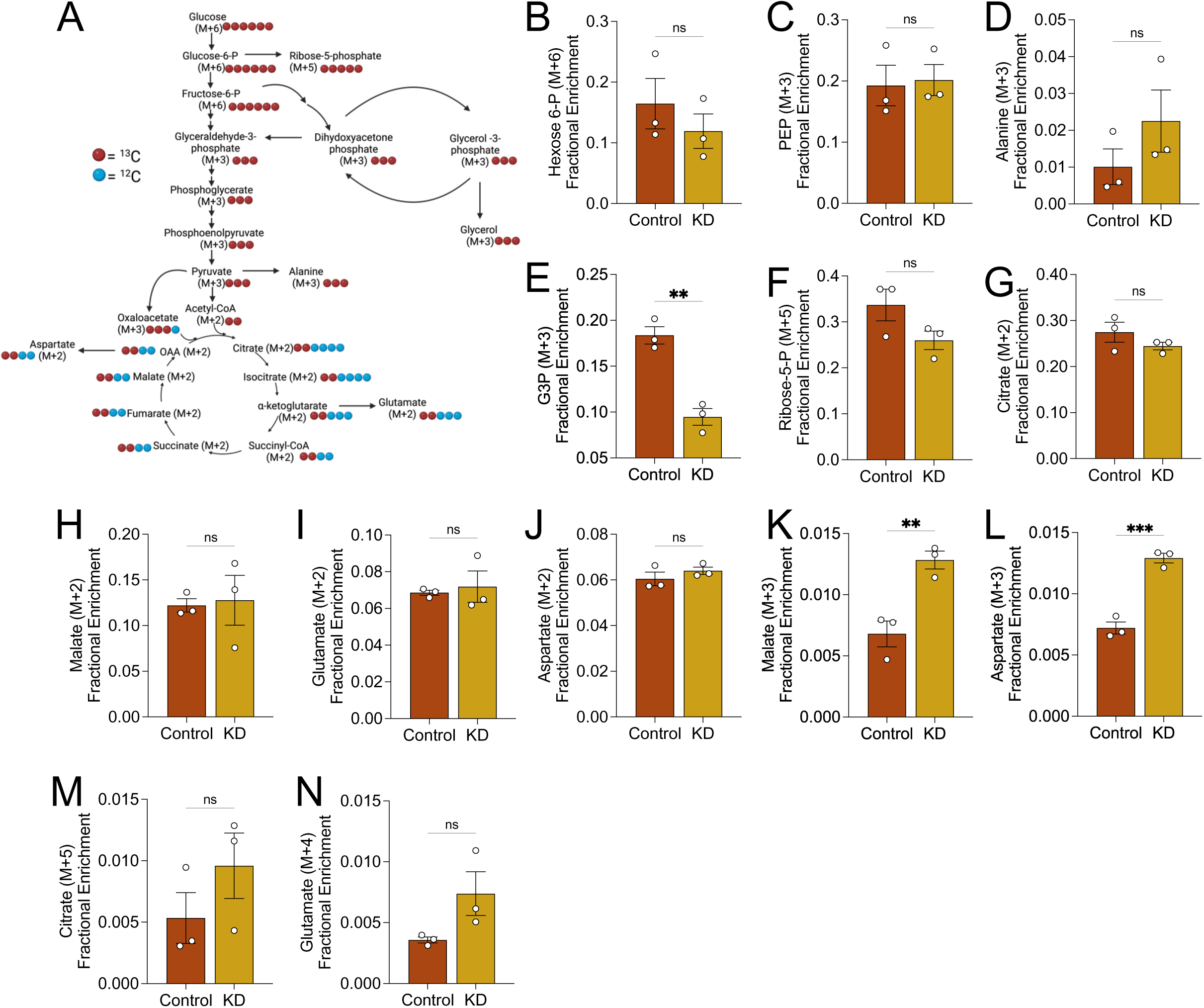
[U-^13^C_6_] glucose Labeling. **(A)** Schematic depicting ^13^C-labelling through glycolysis and the citric acid cycle. Fractional enrichment of **(B)** hexose-6-phosphate (M+6), **(C)** phosphoenolpyruvate (M+3), **(D)** alanine (M+3), **(E)** glycerol-3-phosphate (M+3), **(F)** ribose-5-phosphate (M+5), **(G)** citrate (M+2), **(H)** malate (M+2), **(I)** glutamate (M+2), **(J)** aspartate (M+2), **(K)** malate (M+3), **(L)** aspartate (M+3), **(M)** citrate (M+5) and **(N)** glutamate (M+4) was assessed in cells following incubation with 5.5 mM of [U-^13^C_6_] glucose for 1 h in serum-free culture medium. Data are shown as mean ± SEM and were analyzed by Two-tailed t-test. ^∗^ p< 0.05; ^∗∗^ p< 0.01; ^∗∗∗^ p< 0.001; ∗∗∗∗ p<0.0001.

An important role of glycolysis is to provide intermediates for other linked pathways. Although we were unable to confidently annotate glyceraldehyde-3-phosphate and dihydroxyacetone phosphate in our dataset, we observed a lowered fractional enrichment of M+3 sn-glycerol 3-phosphate, a precursor for triacylglycerols, in KD cells (Fig 3E). We attempted to look at ^13^C incorporation into palmitate and the glycerol backbone of triacylglycerols to determine if glucose-derived carbon units may be incorporated into lipids but found no significant incorporation in palmitate, triacylglycerols or glycerol. We next looked at the ratios of the unlabelled glycerol to sn-glycerol 3-phosphate but found no significant differences between control and knockdown cells (Fig S2K), suggesting no difference in the dephosphorylation to glycerol. Additionally, we observed no differences in the fractional enrichment of M+5 ribose 5-phosphate, a downstream intermediate of the pentose phosphate pathway suggesting no difference in the movement of glucose-derived carbon through the pentose phosphate pathway (Fig 3F).

Glucose-derived carbon units flux through the TCA cycle to yield reducing cofactors and ATP. As mentioned earlier, we were unable to reliably detect pyruvate and acetyl-CoA in our spectra due to noise levels beyond the quantification threshold. However, M+3 pyruvate is oxidatively decarboxylated to M+2 acetyl-CoA, which in turn condenses with oxaloacetate to yield M+2 citrate (Fig 3A). So, fractional enrichment of M+2 citrate can be indicative of glucose oxidation. We detected similar fractional enrichment of M+2 citrate across cell lines (Fig 3G). Further, we observed identical fractional enrichments of M+2 malate, M+2 glutamate, and M+2 aspartate in both cell lines suggesting that the glucose-derived carbon units are fluxed similarly through the TCA cycle (Fig 3H-J).

In addition to flux through acetyl-CoA, pyruvate can be either carboxylated to malate by the malic enzyme or to oxaloacetate by pyruvate carboxylase. Pyruvate flux through either of these alternative pathways will be revealed by a fractional enrichment of M+3 malate or oxaloacetate. We measured the enrichment of aspartate since we were unable to detect oxaloacetate, and oxaloacetate is readily transaminated to aspartate. We found that although the M+3 malate/aspartate isotopologues were trace, the fractional enrichment of M+3 malate and aspartate was elevated in KD cells (Fig 3K-L). Interestingly, our proteomics data revealed an upregulation of pyruvate carboxylase in KD cells, providing support for increased pyruvate carboxylase activity (Fig 6B). We next asked if the carbon units from this alternative pyruvate flux were cycled through the TCA cycle to replenish intermediates. We measured the enrichment of M+5 citrate, which can be derived from the condensation of M+3 oxaloacetate with M+2 acetyl-CoA. We found that KD cells showed elevated enrichment of M+5 citrate although this was not statistically significant (Fig 3M). Interestingly, we observed the same effect on M+4 glutamate (Fig 3N), following the oxidative decarboxylation of M+5 isocitrate to M+4 alpha-ketoglutarate and subsequent transamination of M+4 alpha-ketoglutarate to M+4 glutamate. Taken together, these findings suggest that overall, glucose oxidation is unperturbed in these cardiomyocytes but there is increased flux of pyruvate through malate/oxaloacetate axis in KD cells to potentially replenish citric acid cycle.

### PDK1 suppression decreases mitochondrial fatty acid utilization

Pyruvate dehydrogenase kinases regulation of PDH activity and in turn glucose oxidation, has consequences on fatty acid oxidation. Notably, PDK4 has been shown to increase fatty acid oxidation in the fasted state (37, 38). Therefore, we questioned how fatty acid metabolism is impacted in PDK1 suppressed H9c2 cells. We measured the effect of PDK1 suppression on fatty acid dependency and capacity on palmitic acid treated cells to determine impact on mitochondrial fatty acid utilization. We observed an increase in oxygen consumption rate in the palmitic acid treated cells relative to BSA (Fig 4A). Fatty acid oxidation dependency was lowered in KD cells relative to control cells in both BSA and palmitic acid treated conditions (Fig 4B). We expected an increase in fatty acid oxidation capacity in the palmitic acid treated condition relative to BSA, following the inhibition of alternative pathways such as amino acid and glucose oxidation. Surprisingly, fatty acid oxidation capacity was decreased across cell lines in the palmitic acid treated condition relative to BSA (Fig 4C&D). Further, KD cells showed decreased capacity relative to control only in the BSA condition with no significant differences between cell lines in the palmitic acid-fed condition. These findings suggest that excess palmitic acid perturbs the mitochondrial capacity for fatty acid oxidation in these cells and PDK1 suppression decreases mitochondrial fatty acid utilization independent of nutrient status.

**Figure 4:**
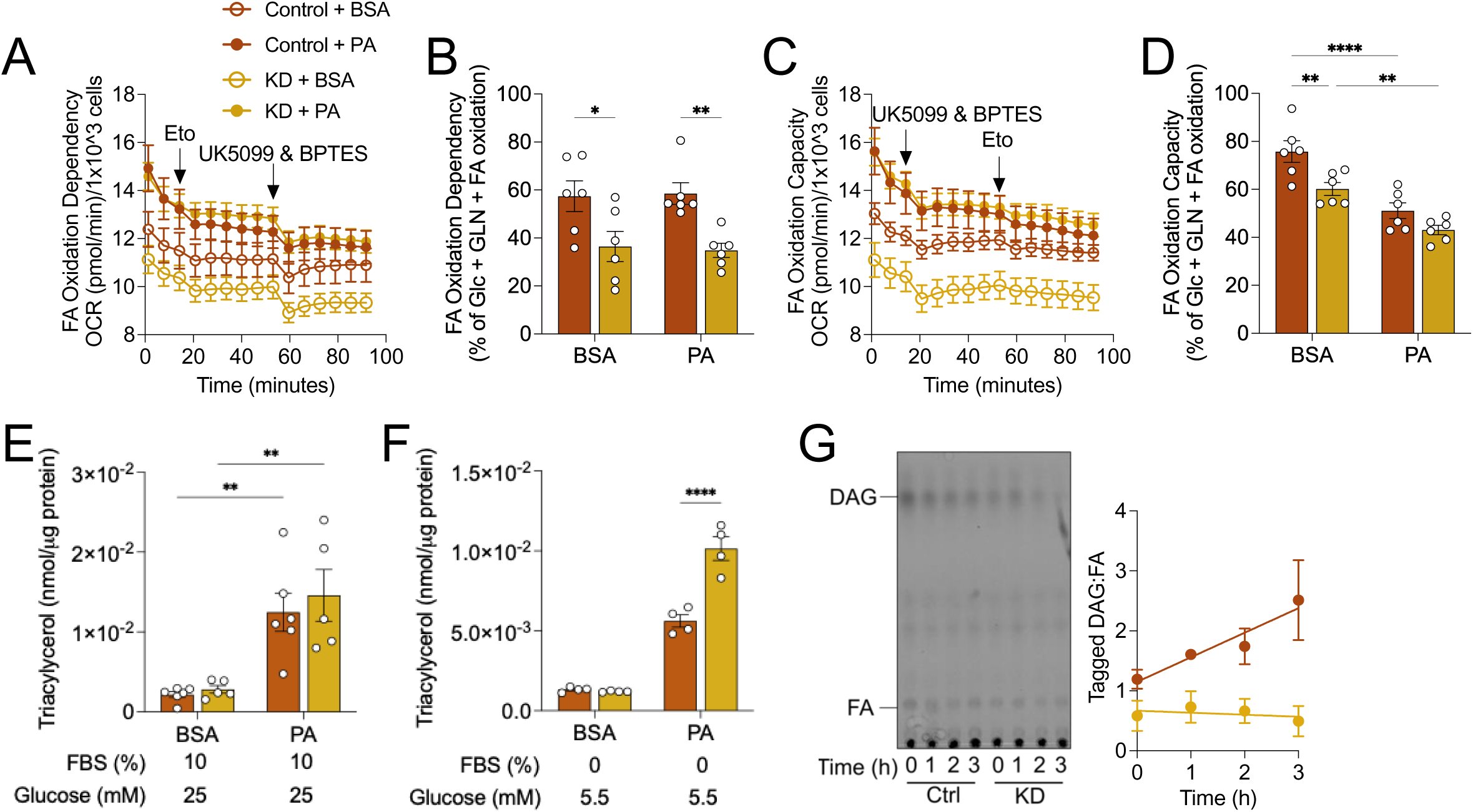
Effect of PDK1 suppression on triacylglycerol metabolism. Cells were incubated in 375 μM of palmitic acid conjugated to BSA (6:1 molar ratio) in complete culture medium for 18 h. **(A&B)** Fatty acid oxidation dependency and **(C&D)** capacity were measured from oxygen consumption rate measurements of 6 independent co-cultures spanning 3 experimental repeats. **(E)** Triacylglycerol abundance was measured following BSA or palmitic acid incubation in complete culture medium, **(F)** nutrient deprived in 5.5 mM glucose, serum free culture media for 3 h. **(G)** Thin layer chromatograph of alkyne incorporated lipids in cells treated with 10 μM of alkyne palmitate for 18 h, then nutrient deprived in low glucose, serum-free media for 3 h (n = 3). Data are shown as mean ± SEM and were analyzed by a Two-way ANOVA followed by a post-hoc Tukey multiple comparison test. For (B), ANOVA showed (F and p values) of (0.066 and 0.7995), (0.002534 and 0.9604), (18.12 and 0.0004) for interaction, row (PA treatment) and column (cell lines) factors respectively. For (D), ANOVA showed (F and p values) of (1.326 and 0.2631), (40.91 and <0.0001), (13.16 and 0.0017) for interaction, row (PA treatment) and column (cell lines) factors respectively. For (E), ANOVA showed (F and p values) of (0.1336 and 0.7190), (30.66 and <0.0001), (0.4757 and 0.4992) for interaction, row (PA treatment) and column (cell lines) factors respectively. For (F), ANOVA showed (F and p values) of (30.91 and 0.0001), (250.2 and <0.0001), (27.78 and 0.0002) for interaction, row (nutrient availability) and column (cell lines) factors respectively. ^∗^ p< 0.05; ^∗∗^ p< 0.01; ^∗∗∗^ p< 0.001; ∗∗∗∗ p<0.0001.

### PDK1 suppression decreases triacylglycerol turnover during nutrient deprivation

We measured triacylglycerol abundance to determine if PDK1 suppression dysregulated triacylglycerol accumulation following excess fatty acid treatment. We found an upregulation of triacylglycerol in both control and KD cells in the palmitic acid-fed condition relative to BSA (Fig 4E). However, we did not observe any differences between cell lines in the basal or palmitic acid-stimulated triacylglycerol abundance (Fig 4E). Since, PDKs generally increase mitochondrial fatty acid utilization during fasting, we turned our attention to the fasted state where triacylglycerols are increasingly broken down to release fatty acids for mitochondrial oxidation (38). Increased mitochondrial fatty acid utilization should increase triacylglycerol breakdown and result in decreased triacylglycerol abundance. So, we fed cells with palmitic acid for 18 h to stimulate triacylglycerol accumulation then nutrient deprived in 5.5 mM glucose, serum free culture media for 3 h. We observed higher levels of triacylglycerol in KD cells relative to control in the low glucose, serum deprived condition (Fig 4F). Pre-treatment of cells with alkyne-palmitic acid to tag the intercellular glycerolipid pool followed by incubation in serum-deprived medium for 3 h revealed increasing diacylglycerol to fatty acid ratios with time in control cells, possibly caused by either increased diacylglycerol production or decreased fatty acid abundance, which could both suggest increased fatty acid utilization (Fig 4G). In contrast, KD cells maintained diacylglycerol-fatty acid ratios across time of serum-deprived fasting (Fig 4G). These findings suggest that PDK1 suppression may negatively impact triacylglycerol turnover during acute mild starvation.

### PDK1 suppression decreases triacylglycerol hydrolysis

To better understand how the lipidome is impacted by PDK1, we extracted lipids from cells fed palmitic acid and nutrient deprived in low glucose medium for 1, 2, and 3 h. Lipids were analyzed using LC-MS/MS and normalized using internal standards spiked into each sample and total protein. We annotated a total of 447 lipid features with MS/MS fragmentation patterns representing 13 lipid classes with peak intensities five times the blank. Since our experiment consisted of three independent variables – cell lines, treatment, and time, we tested which variable was most responsible for the differences in our data. Principal component analysis revealed the formation of two clusters corresponding to the cell lines, along the first principal component responsible for 52.31% of the variance, suggesting that PDK1 knockdown was most responsible for differences in our data (Fig 5A, S3A). We carried out a lipid composition analysis of cells fed with palmitic acid before nutrient deprivation (Time 0) to understand how PDK1 suppression may impact the relative composition of lipid classes. Glycerophospholipids (GPL) were the most abundant lipid category across cell lines in our dataset (Fig 5B&C). Notably, ether-phosphatidylcholine had the highest relative (to total lipid abundance) composition in control while phosphatidylcholine had the highest relative abundance in KD cells (Fig 5B&C), although there was no significant difference in the absolute abundance between these lipid classes between cell lines (Fig 5D). Intriguingly, the relative composition of cholesterol esters was lowered in KD cells and the absolute abundance was significantly downregulated (Fig 5D). The relative composition of glycerolipids (GL) was similar across cell lines with KD cells having slightly higher triacylglycerol and lower diacylglycerol relative compositions (Fig 5B&C). Indeed, there was no significant difference across cell lines in the absolute abundance of the glycerolipids (Fig 5D). We next examined the lipid species that make up the triacylglycerol class. Of the identified triacylglycerol species, triacylglycerol 48:0 (16:0_16:0_16:0) was the most abundant in both cell lines (Fig 5E). Further, there was no statistical difference between cell lines in the absolute abundance across all triacylglycerol species (Fig 5E). These findings reveal only modest differences in lipid composition between control and PDK1-suppressed cells. More importantly, it confirms our findings that PDK1 knockdown does not alter palmitic acid-stimulated triacylglycerol accumulation. We carried out analysis on the lipid acyl chains to determine if PDK1 may be involved in chain modification without changing overall lipid composition. Chain length analysis across all lipid classes revealed no differences between control and KD cells (Fig S3B). However, specific analysis of triacylglycerol chain length reveals lowered abundance of 14C acyl chains and elevated abundance of 17C and 20C acyl chains in KD cells (Fig 5F). Further, triacylglycerol chain saturation analysis revealed higher abundance of acyl chains with two and four double bonds in KD cells (Fig 5G), which was not unique to the triacylglycerol class (Fig S3C). These findings suggest a potential preference for longer and more unsaturated fatty acyl chains in accumulated triacylglycerol in PDK1 knockdown cells. We have previously shown that acute mild starvation modelled by low glucose medium and serum deprivation induces triacylglycerol turnover in control H9c2 cells. So, we compared the lipid abundances across time between control and KD cells to determine the rate of change. We found that triacylglycerol had a significantly lowered rate of disappearance in KD cells, confirming our finding of inhibited turnover in KD cells (Fig 5H&I).

**Figure 5:**
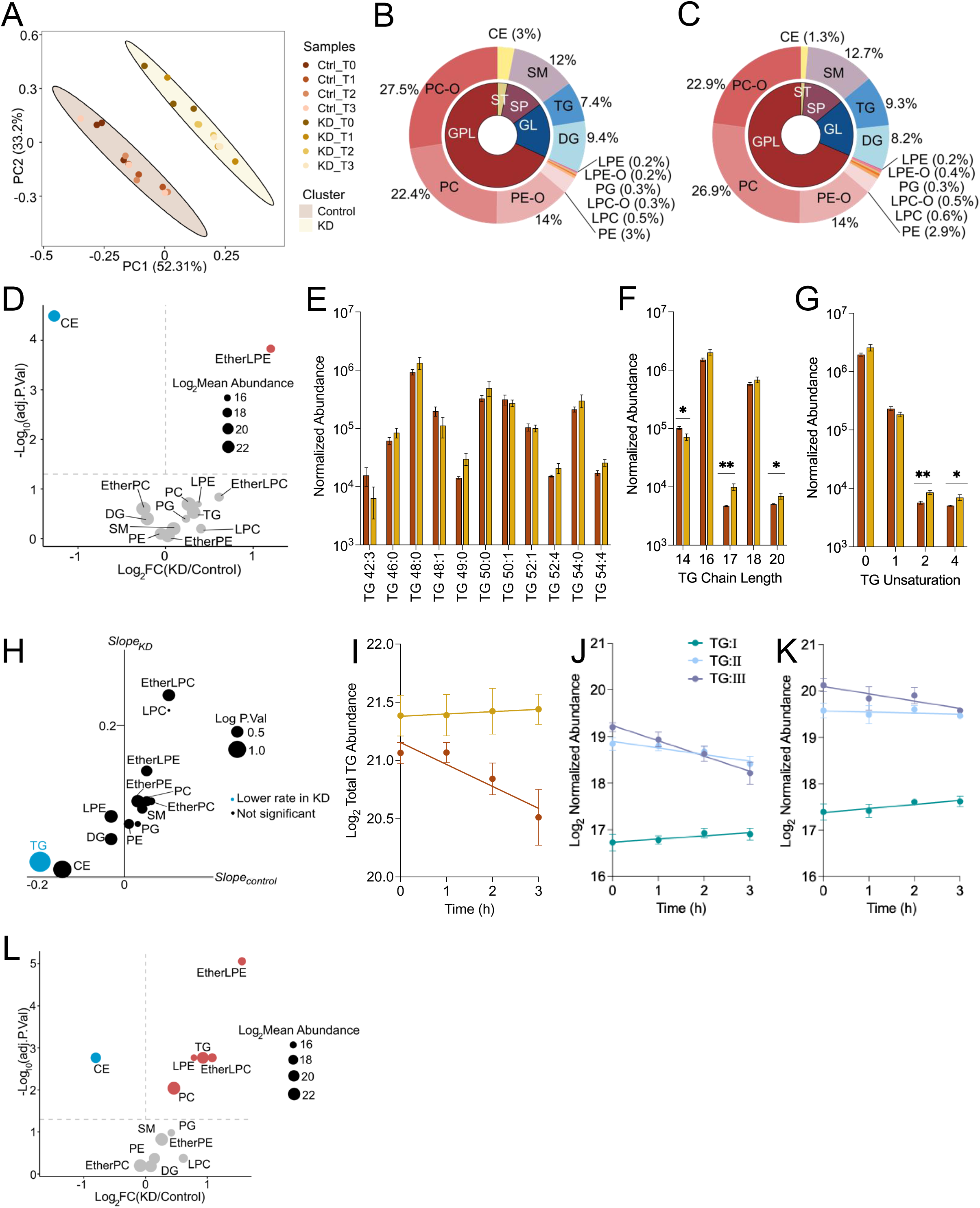
Comparative analyses of lipid profiles. **(A)** Principal component analysis of lipid abundances in control and *Pdk1* knockdown cells treated with 375 μM palmitic acid for 18 h and nutrient deprived in low glucose, serum-free media for 3 h. Relative lipid composition in **(B)** control and **(C)** *Pdk1* knockdown cells following palmitic acid treatment. **(D)** Volcano plot depicting lipid abundances in *Pdk1* knockdown cells relative to control following palmitic acid treatment. Blue and red dots indicate downregulation and upregulation in *Pdk1* knockdown cells respectively. **(E)** Normalized abundance of all identified triacylglycerol species in cells following palmitic acid treatment. Sum abundance of triacylglycerol species of certain **(F)** chain length and **(G)** unsaturation in cells following palmitic acid treatment. **(H)** Slope comparisons of lipid classes in control and *Pdk1* knockdown cells following nutrient deprivation for 3 h. **(I)** Total triacylglycerol abundance during nutrient deprivation, with time = 0 representing the condition in the immediate aftermath of palmitic acid treatment without nutrient deprivation. Cells were treated with 150 μM [U-^13^C_16_] palmitate, then incubated in low glucose, serum-free media for 3 h. Triacylglycerol species with uniformly labeled acyl chains were identified in **(J)** control and **(K)** *Pdk1* knockdown cells, and the sum abundance was plotted against time. N = 3. Data are shown as mean ± SEM and were analyzed by Limma and linear models. ^∗^ p< 0.05; ^∗∗^ p< 0.01; ^∗∗∗^ p< 0.001; ∗∗∗∗ p<0.0001.

Triacylglycerol cycling is a futile cycle that is characterized by the partial hydrolysis and re-esterification of triacylglycerol. Increased triacylglycerol cycling may perturb triacylglycerol turnover. In a labeled experimental paradigm, triacylglycerol cycling would result in the reshuffling of triacylglycerol acyl chains as there is no net loss of acyl chains (39). To better understand triacylglycerol dynamics, we treated cells with [U-^13^C_16_] palmitate and nutrient deprived for 3 h. We resolved ^13^C-labelled triacylglycerols into species with I vs II vs III uniformly labeled acyl chains (ULACs). In control cells, triacylglycerols with 3 ULACs (TG:III) showed the steepest decrease followed by TG:II, while TG:I remained constant (Fig 5J). Summarily, the disappearance of TG:III was not accompanied by increased appearance of triacylglycerols with unlabeled acyl chains, suggestive of no acyl chain shuffling and increased triacylglycerol hydrolysis expected during nutrient deprivation. Interestingly, KD cells show a similar ULAC pattern as in control, with the important distinction of a decreased slope of TG:III, indicating that dysfunctional triacylglycerol hydrolysis, not elevated triacylglycerol cycling, is responsible for lowered rates of change of triacylglycerol levels in KD cells during nutrient deprivation (Fig 5K). Further comparison of lipid class absolute abundance after nutrient deprivation revealed elevated triacylglycerol in KD cells with cholesterol esters still lowered (Fig 5L). These findings strongly suggest that PDK1 knockdown inhibits triacylglycerol hydrolysis in cardiomyocytes.

We performed LC-MS/MS proteomics to map changes in protein abundances that can explain our metabolic phenotype. Principal component analysis showed two main clusters corresponding to cell lines along the first principal component responsible for 30.41% of the variance in the data, indicating that PDK1 knockdown was most responsible for the protein-level differences (Fig S4A). We identified 1861 differentially abundant proteins following palmitic acid treatment and 981 differentially abundant proteins in the nutrient-deprived condition pointing to less proteome dissimilarity in the nutrient-deprived condition (Fig S4B-D). Gene-set enrichment analysis showed positive enrichment of pyruvate and lipid metabolism pathways in nutrient deprived KD cells relative to control cells (Fig S4E). We mapped the protein-protein interaction of the top 150 differentially expressed proteins with the highest significance (Fig S5) and differentially expressed proteins involved in pyruvate and lipid metabolism to their subcellular location (Fig 6) using the Search Tool for the Retrieval of Interacting Genes/Proteins (STRING) database. Despite the decrease in mitochondrial fatty acid utilization, there was significant upregulation of fatty acid oxidation proteins such as carnitine palmitoyltransferase 1a/2 (CPT1a/CPT2), 2,4-dienoyl-CoA reductase 1 (DECR1), acyl-CoA thioesterase 2 (ACOT2), hydroxyacyl-CoA dehydrogenase (HADH), and acyl-CoA dehydrogenase family member 10 (ACAD10) in palmitic acid treated KD cells (Fig 6A). Despite the excess abundance of palmitic acid in the culture medium, we also observed an upregulation of mitochondrial fatty acid synthesis proteins such as malonyl-CoA-Acyl carrier protein transacylase (MCAT) and acyl-CoA synthetase family member 3 (ACSF3) in these cells in both palmitic acid treated and nutrient deprived conditions (Fig 6A&B). These findings hint at proteomic changes that may be adaptive in PDK1 knockdown cells.

**Figure 6:**
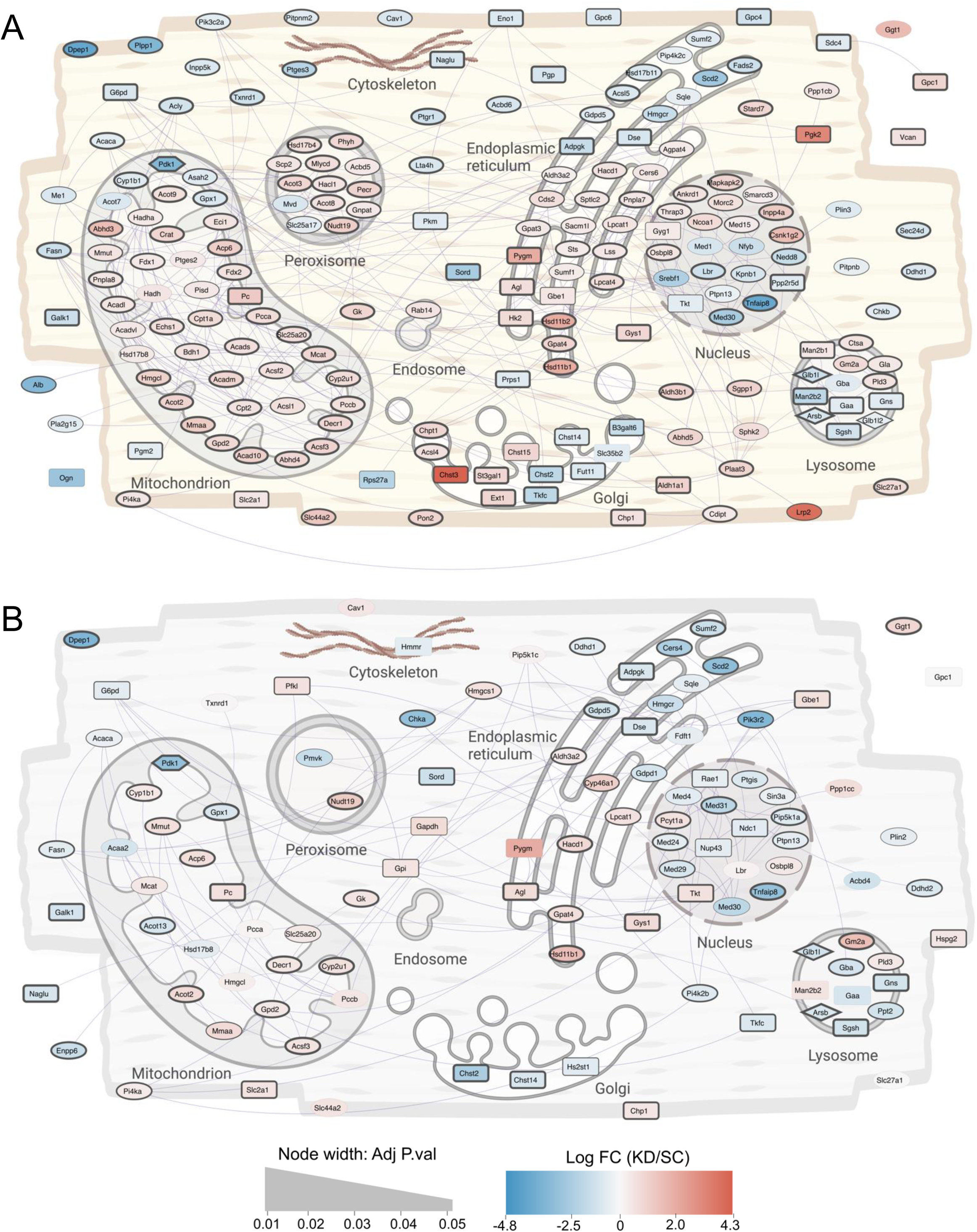
Subcellular mapping of carbohydrate and lipid metabolism proteins. Statistically significant (p < 0.05) differentially abundant metabolism-related proteins between control and *Pdk1* knockdown cells treated with **(A)** 375 μM palmitic acid for 18 h, **(B)** then nutrient deprived in low glucose, serum-free media for 3 h, were connected in a high-confidence (functional score 0.80) protein-protein interaction network (purple lines) using STRING, and depicted in the context of their sub-cellular locations, derived from literature. The color of the nodes represents the fold change in *Pdk1* knockdown cells relative to control, while the thickness of the line around the nodes represents the adjusted p-value. The shape of the node represents the metabolism pathway according to the Reactome database. Circular nodes represent involvement in lipid metabolism, square nodes represent involvement in carbohydrate metabolism, diamond nodes represent involvement in both carbohydrate and lipid metabolism. and the shape of the node represents the metabolism pathway according to Reactome.

A limitation of proteomics data is that it can be challenging to correlate it to metabolic phenotype. Indeed, enzymatic activity can be modulated independent of protein abundance. So, we then used the LipidOne platform (v 2.1) to predict enzymatic activity from our lipidomic dataset (40). Algorithmic prediction revealed decreased activity of adipose triglyceride lipase (ATGL/PNPLA2) in both palmitic acid treated and nutrient deprived KD cells (Fig 7A&B), although there were no changes in ATGL protein abundance coupled with downregulation of perilipin-2 in nutrient deprived KD cells (Fig 7B). Together, these analyses strongly suggest that PDK1 may play a crucial role in regulating triacylglycerol hydrolysis in nutrient deprived cardiomyocytes.

**Figure 7:**
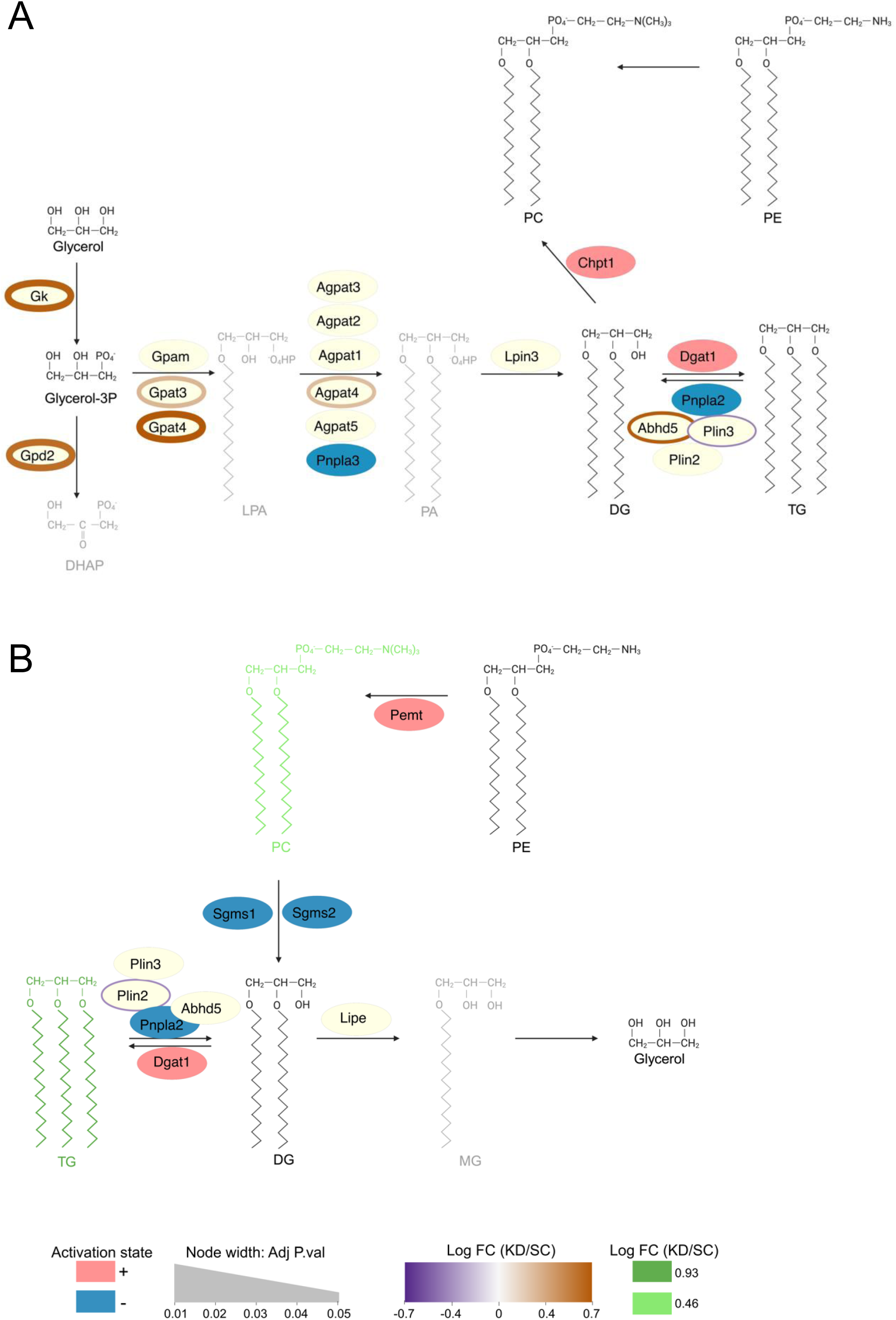
Predicted enzymatic activity. Algorithmic prediction of enzyme activity based on our lipidomics dataset was performed on the LipidOne platform (v 2.1). Activity prediction was carried out in *Pdk1* knockdown cells relative to control in the **(A)** palmitic acid treated condition and the **(B)** nutrient deprived condition. The color of the nodes represents the activation status in in *Pdk1* knockdown cells relative to control. Nodes in cream represent proteins not identified by the algorithm but shown nonetheless to provide increase biochemical context. The color of the line around the node represents the fold change of the protein abundance in *Pdk1* knockdown cells relative to control derived from our proteomics dataset. The thickness of the line around the node represents the adjusted p-value.

### Global PDK1 ablation results in elevated cardiac triacylglycerol levels in fasted male mice

Finally, we tested if the effects of PDK1 on triacylglycerol metabolism in H9c2 cardiomyocytes could be observed in hearts from male global *Pdk1* knockout weaned onto a high fat diet (31) and studied under fasting conditions (Fig 8A). We observed no difference in the body weight and length between WT, HET and KO mice, indicating that PDK1 did not impact overall adiposity (Fig 8B&C). Further, KO mice showed no change in liver, plasma and heart cholesterol (Fig 8D-F). KO mice showed a decrease in plasma triacylglycerol with no changes in liver triacylglycerol abundance (Fig 8G&H). Remarkably, at 12 weeks of age, cardiac triacylglycerol abundance was elevated in 4-hour fasted KO mice relative to WT (Fig 8I), suggesting dysregulated cardiac triacylglycerol metabolism in the absence of PDK1. These data support the *in vitro* findings in our Pdk1 knockdown H9c2 cardiomyocyte model and confirm that PDK1 is a critical regulator of triacylglycerol turnover in the heart.

**Figure 8:**
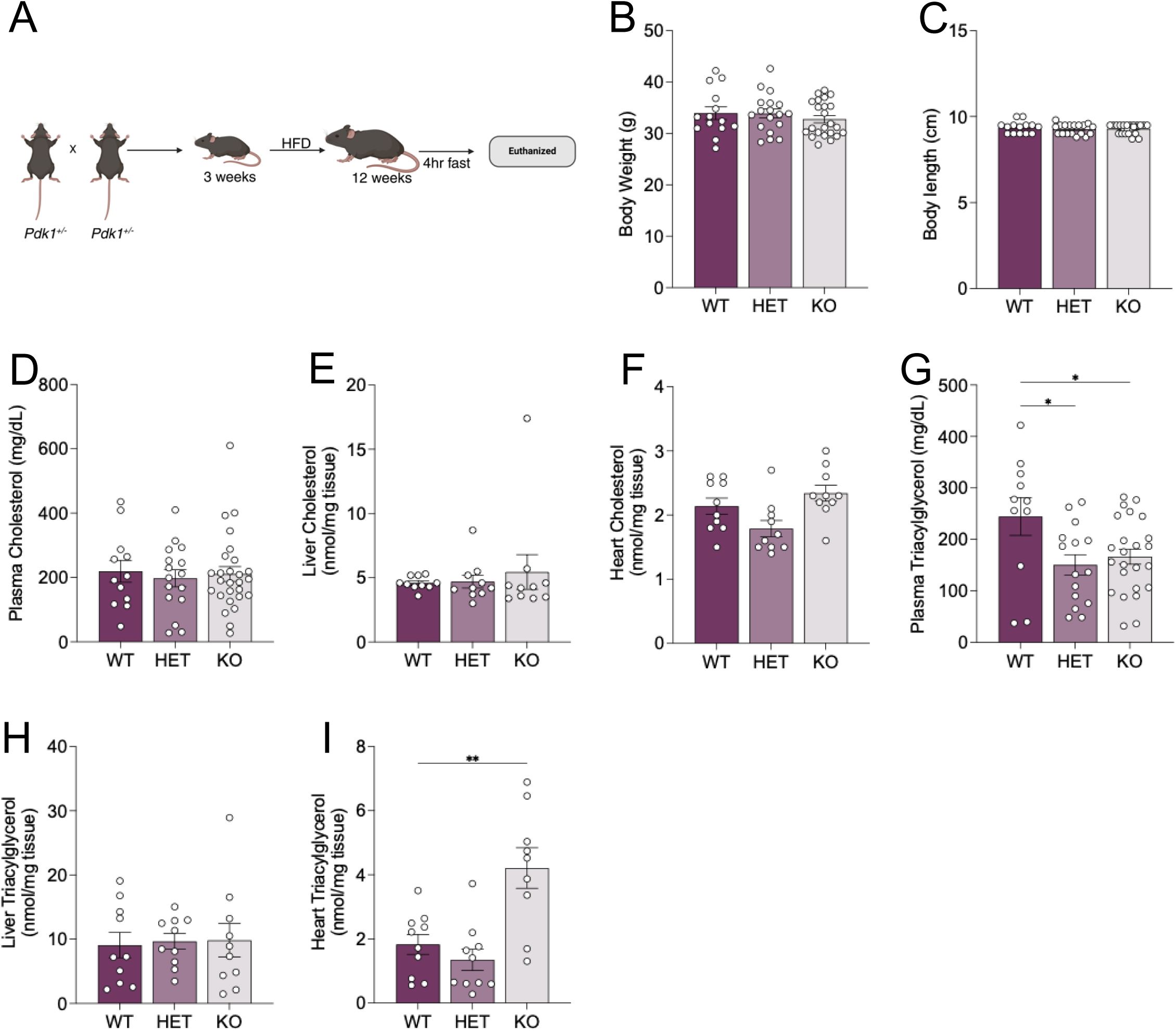
Effect of PDK1 reduction on cardiac triacylglycerol abundance. **(A)** Experimental schematic of 12 weeks aged male mice previously weaned unto a high fat diet were fasted for 4 h. **(B)** Body weight, **(C)** body length, **(D)** plasma cholesterol, **(E)** liver cholesterol, **(F)** heart cholesterol, **(G)** plasma triacylglycerol, **(H)** liver triacylglycerol, and **(I)** heart triacylglycerol was measured. Data are shown as mean ± SEM and were analyzed by One-way ANOVA followed by a post-hoc Dunnett’s multiple comparison test. For (B), ANOVA showed (F and p values) of (0.5775 and 0.5648). For (C), ANOVA showed (F and p values) of (0.4979 and 0.6117). For (D), ANOVA showed (F and p values) of (0.1265 and 0.8814). For (E), ANOVA showed (F and p values) of (0.3098 and 0.7362). For (F), ANOVA showed (F and p values) of (4.986 and 0.0144). For (G), ANOVA showed (F and p values) of (4.228 and 0.0205). For (H), ANOVA showed (F and p values) of (0.03769 and 0.9631). For (I), ANOVA showed (F and p values) of (11.96 and 0.0002), ^∗^ p< 0.05; ^∗∗^ p< 0.01; ^∗∗∗^ p< 0.001; ∗∗∗∗ p<0.0001.

## Discussion

In this study, we use differentiated *Pdk1* knockdown H9c2 cells and *Pdk1* knockout mice to probe the function of PDK1 in cardiomyocytes. We observed that triacylglycerol turnover is attenuated in H9c2 cells following PDK1 loss, and this is accompanied by a decrease in mitochondrial fatty acid utilization. *In vivo* experiments revealed that PDK1 deletion in mice resulted in elevated cardiac triacylglycerol levels in the fasted state with no concomitant increase in body weight or liver triacylglycerol abundance. Further, PDK1’s effect on triacylglycerol metabolism appeared to be independent of glucose metabolism, showing only modest changes with no overall impact on the rate of ATP production. Previous studies have shown increased overall PDK activity in fasted cardiomyocytes (41–43) and have defined the roles of the other PDK isoforms in regulating cardiomyocyte glucose oxidation (44, 45). Our study defines a distinct cardiomyocyte role for PDK1 in the control triacylglycerol turnover to release fatty acids and increase fatty acid oxidation.

In other tissues, PDK1 is crucial in the hypoxic response. Its expression is increased by HIF-1α, the hypoxic response factor (46). Additionally, cytosolic phosphoglycerate kinase 1 translocates to the mitochondria to phosphorylate PDK1, increasing its activity (47). The consequence of these is an inhibition of pyruvate oxidation and increased reliance on glycolysis. However, the metabolic role of PDK1 in cardiomyocytes remained unclear. Complicating the matter further, other findings had revealed no significant change in total pyruvate dehydrogenase kinase activity in response to PDK1 deletion in mice hearts (45). Indeed, our study confirms this finding in H9c2 cardiomyocytes, revealing that PDK1 suppression does not impact total PDH activity (Fig 1F). Surprisingly, we observed seemingly aberrant expression of PDHa and b subunits in KD cells in the palmitic acid-treated condition, with a return to levels in control cells in the nutrient deprived condition (Fig 1G&H). Despite these changes, total PDH activity was consistently similar between control and KD cells in both treatment conditions. Further, In the Klyuyeva study, it was reported that there were no changes in the abundances of PDK2 and 4 following PDK1 deletion (45). Unfortunately, the effect on the abundance of PDK3 was not reported. Here, we observe increased expression in PDK3 following PDK1 reduction (Fig 1I), which may play a compensatory role in maintaining total kinase activity following PDK1 suppression, while counterbalancing the elevated PDH abundance to maintain total PDH activity in these cells. In concert with total PDH activity being unchanged in PDK1-suppressed cells, we observed no changes to total glucose oxidation (Fig 3G-J), further suggesting that PDK1 may be noncritical for the complete oxidation of glucose in cardiomyocytes.

The finding that M+3 sn-glycerol 3-phosphate is decreased in PDK1 KD cells without an observed increase in ^13^C lipid incorporation is intriguing (Fig 3). One potential explanation could be that glucose derived triacylglycerol synthesis does not occur in these cells. The caveat, however, is that technical challenges could be responsible for the lack of observation of ^13^C incorporation and a failure to detect ^13^C incorporation is not an equivalent to evidence for a biological lack of ^13^C incorporation. So, while it is challenging to assess glucose derived triacylglycerol synthesis in our work, it however, seems clear that the rate of consumption of M+3 sn-glycerol 3-phosphate is increased in PDK1 KD cells. Glycerol 3-phosphate is also involved in the glycerol 3-phosphate shuttle to shuttle electrons from the cytosol to the mitochondria, thereby regenerating NAD^+^ to support glycolysis (48). Mining of our proteomics dataset for glycerol 3-phosphate dehydrogenase 1 and 2 (GPD1 and GPD2), the two essential enzyme components of the glycerol 3-phosphate shuttle (48), revealed that although GPD1 was not significantly different, GPD2 was significantly upregulated in PDK1 KD cells (Fig 6). Increase in GPD2 activity may increase the conversion of glycerol-3-phosphate to dihydroxyacetone phosphate in PDK1 KD cells, potentially impacting cellular redox states. Assessments of cytosolic and mitochondrial redox states revealed changes in only mitochondria redox states, which may be supportive of increased GPD2 activity (Fig S6). The caveat, however, is that those assessments were performed using sub-optimally annotated metabolites with lower confidence scores (Progenesis QI scores <45) and no MS2 spectra acquired. Regardless, the consequence of a potential increase in GPD2 activity on the glycerol 3-phosphate shuttle and the ramifications on electron transport in PDK1 KD cells are exciting questions for further exploration.

Fatty acids, the predominant energy source in cardiomyocytes, are characterized by a carboxylic head group and an aliphatic acyl chain (49). Unesterified fatty acids are activated by acyl-CoA synthase to yield fatty-acyl CoA leading to one of two predominant metabolic fates. Fatty-acyl CoA can either be shuttled to the mitochondria where it is oxidized in a series of reactions to yield acetyl-CoA and electron carriers used to produce ATP. Alternatively, fatty-acyl CoA can be shuttled into the intracellular triacylglycerol pool in lipid droplets. Myocardial triacylglycerol pools undergo synthesis and turnover and must be flexible to adapt to the dynamic metabolic and energy demands. Interestingly, Banke et al. (50) showed that myocardial rates of triacylglycerol turnover far exceed the rates of fatty acid oxidation, suggesting predominant flux through intracellular triacylglycerol pools (51), maintained by partial hydrolysis and continuous re-esterification (39, 52). Our findings reveal that PDK1 loss attenuates the disappearance of triacylglycerol during acute mild nutrient deprivation, accompanied by a decreased dependency on fatty acid oxidation (Fig 4B), suggesting reduced fatty acid availability to mitochondria during fasting in the absence of PDK1. This phenotype may be explained by two likely mechanisms. First, PDK1 may directly promote triacylglycerol hydrolysis, and so reduction of PDK1 in the nutrient deprived state results in attenuated triacylglycerol turnover consequently leading to maintenance of triacylglycerol levels. Alternatively, PDK1 may be crucial for fatty acid oxidation such that in absence of PDK1 in the nutrient deprived state, fatty acids are released from triacylglycerol but attenuation of fatty acid oxidation results in re-esterification of fatty acids to triacylglycerol, consequently maintaining triacylglycerol levels in the nutrient deprived state (34). Our investigations with ^13^C_16_ palmitate tracing revealed only modest acyl chain shuffling and substantial differences in the rate of disappearance of TG:III (Fig 5L&M), indicating that triacylglycerol hydrolysis particularly, was dysregulated and suggesting that PDK1 may directly control triacylglycerol turnover.

Triacylglycerol turnover is mediated by two distinct cellular processes – lipolysis and lipophagy (49). Lipolysis is a catabolic process that requires the direct stimulation of lipases, such as Adipose Triglyceride Lipase (ATGL or *Pnpla2*) which catalyzes the rate limiting conversion of triacylglycerol to diacylglycerol (53). Lipophagy, on the other hand, involves the delivery of lipid droplets to lysosomes for degradation (54, 55). However, the relative contribution of these mechanisms to triacylglycerol turnover in cardiomyocytes is unclear. Rambold et al. showed that lipophagy is induced under mild models of nutrient deprivation such as serum-deprivation for 24 h but not in severe models of nutrient deprivation such as HBSS incubation in fibroblasts (56). Alternatively, Jovičić et al. showed that lipolysis is induced in 24 h serum-deprivation of cancer cell lines (57). Regardless, ATGL has been shown to be required for lipolysis and lipophagy by modulating SIRT1, a known activator of lipophagy (58). Therefore, establishing a crosslink between lipolysis and lipophagy. SIRT1 protein abundance appeared to be upregulated in KD cells during nutrient deprivation but fell just outside the arbitrary cut-off for statistical significance (p = 0.056). In contrast, ATGL abundance was unchanged between control and KD cells but was identified as a candidate enzyme with impaired enzymatic activity by algorithmic prediction from our lipidomic dataset (Fig 7B). Interestingly, perilipin 2 protein abundance was significantly downregulated in KD cells.

Perilipins (PLIN) are proteins found on the membrane surface of lipid droplets that help regulate triacylglycerol synthesis and turnover. There are five PLIN isoforms of the protein that differ in tissue expression and regulatory mechanism. PLIN2 and 5 play more significant roles in non-adipose tissues, with PLIN5 being highly expressed in oxidative tissues such as the heart (59). Generally, PLIN2 and 5 inhibit ATGL activity on lipolysis by reducing its localization to lipid droplets and sequestering CGI-58 (*Abhd5*) away from ATGL respectively. Interestingly, the anti-lipolytic activity of PLIN5 can be reversed to pro-lipolytic activity following phosphorylation at S155 to promote interaction with ATGL (60). This pro-lipolytic activity is further enhanced by the ability of PLIN5 to tether lipid droplets to mitochondria to allow for the direct transfer of fatty acids to mitochondria during nutrient deprivation (61). It remains unknown precisely how PDK1, a mitochondrial matrix protein, may regulate a process occurring in lipid droplets. However, perilipins could be ideal candidates worth further investigation. Specifically, if they are substrates of PDK1 during nutrient-deprivation to promote fatty acid availability to the mitochondria.

A limitation of our study is that it is difficult to ascertain which effects are direct or indirect effects of *Pdk1* knockdown. Although pharmacological inhibition is subject to the same challenge, it could be a useful orthogonal experiment in validating the reported effects of PDK1 suppression. The additional challenge with pharmacological inhibition is a lack of drugs that specifically inhibit PDK1 and not the other PDK isoforms. Secondly, majority of PDK pan-inhibitors act by disrupting the interaction/reaction between PDKs and PDH. Our study suggests that PDK1 effects on triacylglycerol metabolism may be independent of PDH and carbohydrate metabolism. Therefore, a need for specific inhibitors that disrupt PDK1 kinase activity remains. Additionally, our in vivo experiments, performed only in male mice, leave questions about sex differences and the impact of PDK1 deletion on cardiac function, unanswered. These limitations create intriguing questions worthy of exploration in further studies.

## Experimental procedures

### Animals

Animals were housed in ventilated cages within the Modified Barrier Facility at the University of British Columbia with 12-h light:dark cycles. All mice were generated from crosses between heterozygotes (HETs) such that littermates were used as controls. A heterozygous breeder pair of Pdk1^tm1.1(KOMP)Vlcg^ knockout (KO), global knockout mice, were obtained from the Knockout Mouse Project (KOMP), then bred in-house and backcrossed to the C57BL/6J. Because prior studies showed no effect of this locus in females, experiments used exclusively male animals (31). Experimental mice were weaned at 3 weeks on a diet high in fat with 20% kcal sucrose, and 60% kcal fat (D12492i, Research Diets). At 12 weeks, mice were fasted for 4 h and euthanized by an overdose of isoflurane anesthesia in accordance with the Animal Care guidelines. At this time a cardiac blood sample was withdrawn, and tissues were rapidly harvested, weighed, and flash frozen. Blood samples were kept on ice, then plasma was separated by centrifugation at 4°C at 10,000 rpm for 8 min. Tissue and plasma samples were stored at −80°C until analysis. All procedures were approved by the University of British Columbia Animal Care and Use Committee and were performed in accordance with Canadian Council on Animal Care Guidelines (Protocol No. A19-0267).

### Tissue Cholesterol and Triacylglycerol

Metabolite levels were measured in plasma from the cardiac blood samples collected from mice at the time of euthanasia. Plasma cholesterol (Cholesterol-SL Assay, Cat. No. 234-60 Sekisui Diagnostics, PEI, Canada) and triacylglycerol (Triglyceride-SL Assay, Cat. No. 236-60, Sekisui Diagnostics, PEI, Canada) were measured using colorimetric assays according to the manufacturers’ directions.

Tissue lipids were extracted from liver and heart samples as previously described (31). Tissue cholesterol abundance was measured using a colorimetric assay (Cholesterol E, Cat. No. 439-17501, Wako Diagnostics, Richmond, VA) according to the manufacturer’s directions. Tissue triacylglycerol abundance was measured using a colorimetric assay (Glycerol Reagent, Cat. No. F6428, Triglyceride Reagent, Cat. No. T2449, Sigma-Aldrich, Oakville, ON, Canada) according to the manufacturer’s instructions.

### H9c2 Knockdown Cell Line Generation and Differentiation

H9c2 cells (Cat. No. 88092904, Sigma-Aldrich, Oakville, ON, CA) were cultured in complete medium containing high glucose (25 mM) Dulbecco’s Modified Eagle Medium (Cat. No. 30-2002, ATCC, Manassas, Virginia, US) supplemented with 10% fetal bovine serum (Cat. No. 12483020, ThermoFisher Scientific, Burnaby, BC, CA) and 1% antibiotic-antimycotic containing 100 units/mL of ThermoFisher Scientific, Burnaby, BC, CA) in a 37 °C, 5% CO_2_ cell culture incubator. Pdk1 shRNA penicillin, 100 μg/mL of streptomycin and 0.25 μg/mL of Amphotericin B (Cat. No. 15240062, Lentiviral particles (Cat. No. TL712182V, Origene, Rockville, Maryland, US) were mixed with 8 μg/mL of polybrene (Cat. No. TR-1003, Sigma-Aldrich, Oakville, ON, CA) and added to cultured cells at a multiplicity of infection (MOI) of 500 or 120 (for the scrambled shRNA control). Stably transduced cells were positively selected by daily treatment with 1.5 μg/mL of puromycin (Cat. No. P8833, Sigma-Aldrich, Oakville, ON, CA) until cell dish became 70-80% confluent.

H9c2 cells were seeded in a culture flask containing complete media and incubated in the cell culture incubator for 24 h to allows cells attach to the flask. Complete media was replaced with differentiation media containing high glucose (25 mM) Dulbecco’s Modified Eagle Medium supplemented with 1% fetal bovine serum, 1% antibiotic-antimycotic and 1μM *trans* retinoic acid (Cat. No. 554720, Sigma-Aldrich, Oakville, ON, CA). Cells were maintained in fresh differentiation media daily for 10 days.

### Palmitic Acid Treatment, Palmitic acid alkyne and ^13^C Labelling

Differentiated cells were treated with 375 μM palmitic acid complexed with fatty acid free bovine serum albumin in a 6:1 molar ratio for 18 h, after which cells were placed in ‘starvation’ media containing low glucose (5 mM) Dulbecco’s Modified Eagle Medium (Cat. No. 11885084, ThermoFisher Scientific, Burnaby, BC, CA) for 3 h.

Cells were labelled with ^13^C_16_ palmitate by treating with media containing Dulbecco’s Modified Eagle Medium supplemented with 2% fatty acid free bovine serum albumin, 1% antibiotic-antimycotic and 0.15 mM ^13^C_16_ palmitate (Cat. No. 605573, Sigma-Aldrich, Oakville, ON, CA) complexed to bovine serum albumin for 18 h after which they were placed in ‘starvation media’ for 3 h with cells collected every hour. Cells were dynamically labelled with ^13^C_6_ glucose by incubating first in unlabelled ‘starvation media’, then replacing glucose in the starvation media with 5.5 mM of ^13^C_6_ glucose (Cat. No. 26707, Cayman Chemical Company, Michigan, US) for 2, 5, 10, 20, 30, and 60 mins. The total time incubated in starvation media was 3 h in every instance. Cells were labelled with palmitic acid alkyne by treating with media containing Dulbecco’s Modified Eagle Medium supplemented with 10% fatty acid fetal bovine serum, 1% antibiotic-antimycotic and 10 μM of palmitic acid alkyne (Cat. No. 13266, Cayman Chemical Company, Michigan, US) for 18 h after which they were placed in ‘starvation media’ for 3 h with cells collected every hour. After labelling, cells were washed with ice cold phosphate buffered saline (PBS), scraped on ice, centrifuged at max speed at 4 °C for 5 mins, and pellet stored at -80 °C.

### Gene Expression Analysis

H9c2 cells were harvested from culture flasks by discarding old media and a 5 min incubation in the cell culture incubator in 0.25% trypsin-EDTA (Cat. No. 25200072, ThermoFisher Scientific, Burnaby, BC, CA). Cell suspension was spun at 300 xg for 5 mins. The cell pellet was washed with PBS and re-spun at 300 xg for 5 mins and the supernatant was discarded. Total RNA was extracted from cell pellets using the Qiagen RNeasy extraction kit (Cat. No. 74104, Qiagen, Hilden, DE), after which cDNA was synthesized using the qScript cDNA synthesis kit (Cat. No. 101414-100, Quanta Biosciences, Beverley, US). Gene expression was measured with using real time qPCR and Taqman gene expression assay (ThermoFisher Scientific, Burnaby, BC, CA). β-actin (*Actb*, Cat. No. Rn01410374_m1, ThermoFisher Scientific, Burnaby, BC, CA) was used as a house keeping gene. Cycle threshold (Ct) values of genes (*Pdk1*, Cat. No. Rn00587598_m1; *Pdk2*, Cat. No. Rn00446679_m1; *Pdk3*, Cat. No. Rn01424337_m1; *Pdk4*, Cat. No. Rn00585577_m1, ThermoFisher Scientific, Burnaby, BC, CA) were subtracted from Ct values of β-actin to obtain relative expression to β-actin (ΔCt). ΔCt values of treated cells were subtracted from ΔCt values of controls to obtain fold change relative to control (ΔΔCt), which was then converted to linear scale (2^ΔΔCt^).

### Western Blotting

Immunoprecipitation assay lysis buffer containing 150 mM NaCl, 1% Nonidet P-40, 0.5% sodium deoxycholate, 0.1% sodium dodecyl sulfate (SDS), 50 mM Tris, pH 7.4, 2 mM EGTA, 2 mM Na_3_VO_4_, and 2 mM NaF supplemented with complete mini protease inhibitor cocktail (Cat. No. 11836170001, Sigma-Aldrich, Oakville, ON, CA) was added to cell pellets, frozen at -80 °C and thawed. Lysates were spun in a microcentrifuge at 12,000 rpm, 4 °C for 10 mins. Protein concentration was measured in the supernatant using a Pierce bicinchoninic acid protein assay kit (Cat. No. 23225, ThermoFisher Scientific, Burnaby, BC, CA). Lysates were incubated in Laemmli loading buffer (Cat. No. J61337AC, ThermoFisher Scientific, Burnaby, BC, CA) containing dithiothreitol at 95 °C for 5 mins. Equal amounts of protein from each sample were resolved on a 4% stacking, 12% running SDS-PAGE. Proteins were transferred to PVDF membranes (Cat. No. 162-0177, Bio-Rad, California, US) and blocked in 5% skim milk/TBS-T for 1 h. Membranes were probed with primary antibodies against PDK1 (1:2 500 in 5% skim milk/TBS-T, Cat. No. ab202468, Abcam, Cambridge, UK), β-tubulin (1:2 000 in 5% skim milk/TBS-T, Cat. No. T8328, Sigma-Aldrich, Oakville, ON, CA). The signals were detected by secondary horseradish peroxidase conjugated antibodies (1:10 000 in 5% skim milk/TBS-T, Anti-mouse, Cat. No. 7076; 1: 10 000 in 5% skim milk/TBS-T, Anti-rabbit, Cat. No. 7074S, Cell Signaling Technology, Massachusetts, US) and Pierce ECL Western Blotting Substrate (Cat. No. 32106, ThermoFisher Scientific, Burnaby, BC, CA).

### PDH Activity Assays

Cells were harvested from culture flasks by scraping in ice cold PBS and centrifuged at 10, 000 xg for 5 mins at 4 °C. Cell pellet was then resuspended in assay buffer from the pyruvate dehydrogenase activity kit (Cat. No. MAK183, Sigma-Aldrich, Oakville, ON, CA). Cell suspension was sonicated at the 45% amplitude with 3 pulses on for 10 secs and off for 10 secs, after which they were kept on ice for 15 mins and centrifuged at 10, 000 xg for 5 mins at 4 °C. Enzyme activity was measured per manufacturer’s instructions.

### Seahorse Assay

Seeded cells at 1.5 x 10^4^ cells per well of a Seahorse XFe96 cell culture in complete growth medium. After letting cells attach for 24 h, cells were treated with 375 μM palmitic acid complexed with fatty acid free bovine serum albumin for 18 h. Oxygen consumption rate and extracellular acidification rate were then measured following the manufacturer’s kit instructions. To calculate % nutrient dependency, we used the following formula as recommended by the manufacturer:

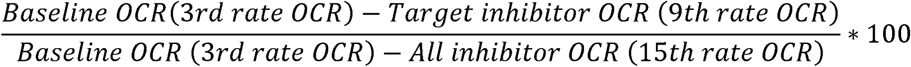

To calculate % nutrient capacity, we used the following formula as recommended by the manufacturer:

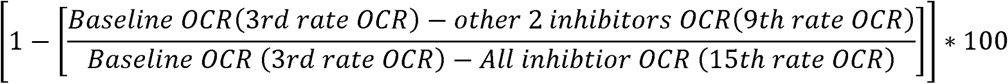

### Triacylglycerol Assay

Cells were harvested from culture flasks by scraping in ice cold PBS and centrifuged at 10, 000 x g for 5 minutes at 4 °C. Cell pellets were resuspended in 5% Nonidet P-40 and sonicated at 45% amplitude with 3 pulses on for 10 secs and off for 10 secs, after which they were incubated at 95 °C for 5 minutes, cooled back down to room temperature and centrifuged at 20, 000 x g for 3 mins. Triacylglycerol abundance was then measured in the supernatant using a triacylglycerol assay kit (Cat. No. ab65336, Abcam, Cambridge, UK).

### Click Chemistry and Thin Layer Chromatography

Lipids were extracted and alkyne-modified lipids were reacted with 3-azido-7-hydoxycoumarin (Cat. No. 909513, Sigma-Aldrich, Oakville, ON, CA) and resolved on a TLC plate as previously described (62). Resolved lipids were visualized with the ChemiDoc Imaging System and quantified by performing densitometry using ImageJ.

### Proteomics

Cell pellets were diluted with ultrapure water, then lysed by adding 50% cold trifluoroethanol (TFE). Samples were then chilled on ice for 10 min, vortexed for 1 min and sonicated for 5-10 min in ice bath. One hundred mM Tris (pH 8-8.5) and Tris(2-carboxyethyl)phosphine (1 μg/50 μg protein) was added to the samples, and incubated for 20 min, then alkalized with chloroacetamide (1 μg/10 μg protein), incubated at 95°C for 10 min. Samples were diluted with 50 mM ammonium bicarbonate, digested with LysC/Trypsin (1 μg/50 μg protein) for 2 h at 37°C, and incubated further at 37°C overnight. Samples were further digested with trypsin (1 μg/125 μg protein) for 5 h at 37°C. Digestion was stopped by acidification and samples were desalted with 6 mm C18 stage tips conditioned with 100% methanol, equilibrated and washed with 0.2% trifluoroacetic acid, and finally eluted with 40% acetonitrile, 0.1% formic acid. Eluate was dried and reconstituted in 0.5% acetonitrile, 0.1% formic acid before instrumental analysis.

One hundred ng of peptides were injected and separated on-line using NanoElute UHPLC system (Bruker Daltonics) with Aurora Series Gen2 (CSI) analytical column, (25 cm x 75 μm 1.6 μm FSC C18, with Gen2 nanoZero and CSI fitting; Ion Opticks, Parkville, Victoria, Australia) heated to 50°C and coupled to timsTOF Pro (Bruker Daltonics). A standard 30 min gradient was run from 2% B to 12% B over 15 min, then to 33% B from 15 to 30 min, then to 95% B over 0.5 min, and held at 95% B for 7.72 min. Before each run, the analytical column was conditioned with 4 column volumes of 0.1% aqueous formic acid and 0.5 % acetonitrile in water, and buffer B consisting of 0.1% formic acid in 99.4% acetonitrile. The NanoElute thermostat temperature was set at 7°C and the analysis was performed at 0.3 μL/min flow rate.

The Trapped Ion Mobility – Time of Flight Mass Spectrometer (TimsTOF Pro; Bruker Daltonics, Germany) was set to Parallel Accumulation-Serial Fragmentation (PASEF) scan mode for DIA acquisition scanning 100 – 1700 m/z. The capillary voltage was set to 1800 V, drying gas to 3 L/min, and drying temperature to 180°C. The MS1 scan was followed by 17 consecutive PASEF ramps containing 22 non-overlapping 35 m/z isolation windows, covering the m/z range 319.5 – 1089.5. As for the TimsTOF setting, ion mobility range (1/k0) was set to 0.70 – 1.35 V·s/cm^2^, 100ms ramp time and accumulation time (100% duty cycle), and ramp rate of 9.42Hz, resulting in 1.91s of total cycle time. The collision energy was ramped linearly as a function of mobility from 27eV at 1/k0 = 0.7 V·s/cm^2^ to 55eV at 1/k0 = 1.35 V·s/cm^2^.

Mass accuracy: error of mass measurement was not allowed to exceed 7 ppm. The TimsTOF Pro instrument was run with timsControl v. 4.1.12 (Bruker). LC and MS were controlled with HyStar 6.0 (6.2.1.13, Bruker). Acquired diaPASEF data were then searched using DIA-NN (63) (v. 1.8.1) to obtain DIA quantification, with the spectral library generated from the canonical proteome for Rattus Norvegicus from Uniprot. Quantification mode was set to “Any LC (high precision)” and a two-pass search was completed. All other settings were left default.

### Cytoscape Processing

The initial proteomics datasets were first filtered to exclude features mapped to multiple possible genes, and all features with the term ‘BOVIN’ in the protein group as these were likely residual contaminants from FBS used in culture. To identify proteins from our proteomics data that are involved in either carbohydrate or lipid metabolism, Reactome (https://reactome.org/) was used to generate lists of proteins from *Rattus norvegicus* under the categories “Metabolism of lipids” and “Metabolism of carbohydrates”. The resulting uniprot IDs were then converted to gene names using the Uniprot Retrieve/ID mapping tool (https://www.uniprot.org/id-mapping). The proteomics dataset was queried for a match. High-confidence (STRING functional score 0.8) protein-protein interaction network was generated for all statistically significant (adjusted p < 0.05) metabolism-related differentially expressed proteins between the control and PDK1 knockdown cells treated in palmitic acid treated and nutrient-deprived conditions. PDK1 was not a result in the Reactome metabolism results but was plotted in a distinct shape. Subcellular location was assigned as the location with the highest score (0-5) assigned in STRING. When there was a tie, PubMed was queried to determine the most appropriate location. A medium-confidence (STRING functional score 0.5) protein-protein interaction was also generated for the top 150 (smallest p-value) differentially expressed proteins in each condition. The same methods were followed as for the metabolism networks, but proteins that were in the top 150 of both conditions were highlighted with a green node border to illustrate similarities.

### Lipidomics LC-MS/MS Analysis

Lipidomic analysis was conducted following previously established protocols with some modifications (64, 65). Before LC−MS/MS analysis, lipid extracts were resuspended in 200 µL of an acetonitrile solution: isopropanol (7:3, v/v) containing 100 ng/mL of CUDA and 3 µl of SPLASH® LIPIDOMIX® Mass Spec Standard as an internal standard. The samples were vortexed for 10 minutes and subsequently centrifuged at 14,000 rpm for 10 minutes. 150 µL were transferred to LC vials for lipidomics analysis. Aliquots of 20 µL from each sample were pooled to generate a quality control (QC) sample, which was used to monitor LC-MS/MS platform performance.

Untargeted lipidomics analysis was performed using an Impact II™ high-resolution mass spectrometer (Bruker Daltonics, Bremen, Germany) coupled with a Vanquish UHPLC system (Thermo Fisher Scientific, Waltham, MA). Separation of lipid species was achieved on an Acquity CSH (Charged Surface Hybrid) C18 column (130Å, 1.7 µm, 100 x 2.1 mm) (Waters, Milford, MA) equipped with a CSH C18 VanGuard FIT Cartridge (1.7 µm, 2.1 x 5 mm), utilizing a multigradient adapted from Cajka et al., (64). The mobile phase A consisted of acetonitrile (60:40, v/v), while mobile phase B comprised isopropanol (90:10, v/v). Both phases contained 0.1% (v/v) formic acid and 10 mM ammonium formate. Separation was executed using a gradient ranging from 15% to 99% mobile phase B over 17 minutes as follows: 0 min, 15% B; 0–2 min, 30% B; 2–2.5 min, 50% B; 2.5–12 min, 80% B; 12–12.5 min, 99% B; 12.5–13.5 min, 99% B; 13.5–13.7 min, 15% B; 13.7-17 min, 15% B. The column temperature was maintained at 65°C, with a flow rate of 0.5 mL/min. The injection volume was 2 µL, and the autosampler temperature was kept at 4°C.

Data was recorded in the data-dependent acquisitions, with both positive (ESI+) and negative (ESI-) ionization modes used to obtain precursor and fragment ion data for compound annotation. For ESI+, the mass spectrometer settings were as follows: capillary voltage, 4500 V; nebulizer gas pressure, 2.0 bar; dry gas flow rate, 9 L/min; dry gas temperature, 220°C; mass scan range, 65–1700 *m/z*; spectra acquisition rate, 3 Hz; cycle time, 0.7 s. Collision energy was ramped from 100% to 250% during each MS/MS scan, starting at 20 V. For ESI-, the capillary voltage was set at -4000 V. Calibration was performed by injecting 10 µL of 10 mM sodium formate at the start of each run via the 6-port diverter valve. To assess instrument performance, QC samples were injected every nine samples.

Lipid features with MS/MS were then annotated using MS-DIAL (v 5.1.230912) and involved peak picking, alignment, and LIPID MAPS database searching (66). Lipid classes were normalized by internal standard peak height and protein concentration. Duplicate features were filtered out based standard deviation in the QC samples – duplicate features with the smallest deviation were kept. Additionally, features with the highest sample peak area less than 5x the value of the blank was filtered out and regarded as not meeting the quantification threshold. The dataset was then transformed to log_2_ scale to normalize the data distribution before statistical tests were performed. Additionally, it was also analyzed on the LipidOne (v 2.1) platform to foster biological interpretation (40).

### Metabolomics LC-MS/MS Analysis

Samples were analyzed following previously established protocols with some modifications (67, 68). The polar fraction was resuspended in 200 µL of 50% aqueous methanol (v/v) spiked with internal standards at 1 ppm (methionine-d3, ferulic acid-d3, caffeine-^13^C_3_). Following resuspension, the mixture was centrifuged at 14,000 rcf for 10 minutes, and 150 µL of the resulting supernatant was transferred to LC-MS vials (Thermo Fisher) for subsequent LC-MS/MS analysis. To assess analytical performance, 20 µL aliquots from each sample were pooled to create a quality control (QC) sample.

Metabolite analysis was conducted using an Impact™ II high-resolution mass spectrometer (Bruker Daltonics, Bremen, Germany) and a Vanquish Horizon UHPLC system (Thermo Fisher Scientific, Waltham, MA). Separation of compounds was performed using a multigradient method on an Inertsil Ph-3 UHPLC column (2 µm, 150 x 2.1 mm, GL Sciences), which was equipped with a Ph-3 guard column (2 µm, 2.1 x 10 mm). The mobile phase consisted of water (A) with 0.1% (v/v) formic acid and methanol (B) with 0.1% (v/v) formic acid. Chromatographic separation was achieved using a multi-step gradient from 5% to 99% mobile phase B over 18 minutes. The UHPLC program was set as follows: 0 min (5% B); 0–1 min (5% B); 1–8 min (35% B); 8–10.5 min (99% B); 10.5–14 min (99% B); 14–14.5 min (5% B); 14.5–18 min (5% B). The column temperature was maintained at 55°C, with the autosampler kept at 4°C, and a flow rate of 0.3 mL/min was employed - 5 µL injected in both positive and negative ionization modes.

Data-dependent acquisitions were performed in both positive (ESI+) and negative (ESI-) ionization modes, to capture precursor and fragment ions for compound annotation. For ESI+, the mass spectrometer parameters were as follows: capillary voltage of 4,500 V, nebulizer gas pressure of 2.0 bar, dry gas flow rate of 9 L/min, dry gas temperature of 220°C, mass scan range of 60–1,300 *m/z*, and a total cycle time of 0.6 s. To ensure comprehensive structural information, collision energy of 20 V was ramped from 100% to 250% during each MS/MS scan. For ESI-, the capillary voltage was set at -3,500 V. High mass accuracy was maintained by performing internal calibration during each analytical run using 10 µL of 10 mM sodium formate, injected at the start of the run (from 0 to 0.15 min) via a 6-port valve.

Quality control samples and blanks were systematically injected every nine samples to track instrument performance. To mitigate potential batch effects, samples were fully randomized before injection. Metabolite features in unlabeled samples were annotated and normalized as previously described (67). The retention times and *m/z* values were used in predicting labelled isotopologues and MS1 peaks were identified in Skyline (v 24.1) (69). We identified all possible isotopologues for each metabolite and calculated the fractional enrichment as a ratio of the isotopologue abundance to the sum of all isotopologues for a given metabolite and then carried out a natural abundance correction as described previously (70).

### Statistical Analysis

Statistical analyses and data presentation for were carried out using GraphPad Prism 9 (Graphpad Software, San Diego, CA, USA) or R (v 4.1.1). Two-tailed Student’s *t*-test was performed for pairwise comparisons. For omics data, linear models for microarray (LIMMA) R package (71) was used in assessing differential abundance. For all statistical analyses, differences were considered significant if the (adjusted) p-value was less than 0.05: ^∗^ p< 0.05; ^∗∗^ p< 0.01; ^∗∗∗^ p< 0.001; ∗∗∗∗ p<0.0001. Data were presented as means ± SEM with individual data points from biological replicates, unless otherwise indicated in the figure legends.

## Data availability

The raw data supporting the conclusions of this article will be made available by the authors without reservations. The R scripts are available on GitHub at https://doi.org/10.5281/zenodo.14707488. The proteomics mass spectra have been uploaded unto ProteomeXchange (PXD059927). The lipidomics and metabolomics mass spectra are available at the NIH Common Fund’s National Metabolomics Data Repository (NMDR) website, the Metabolomics Workbench, https://www.metabolomicsworkbench.org where it has been assigned Study IDs ST003688 and ST003689 respectively. The data can be accessed directly via its Project DOI: http://dx.doi.org/10.21228/M8W546 (72).

## Supporting Information

This article contains supporting information.

## Acknowledgments

We acknowledge all the members of the Johnson lab for their supporting efforts in the completion of this study and thank the anonymous referees for their useful suggestions in strengthening our manuscript.

## Funding and additional information

This work was supported by a CIHR Project Grant to Suzanne Clee (PTJ-005633), which was transferred to J.D.J. M.G.A was supported by the University of British Columbia CELL 1Y fellowship. E.M.R was supported by a Vanier Canada Graduate Scholarship. E.D.Q and P.C were supported by the National Institutes of Health (NIH T32HL166142 and NIH R01DK091538, respectively). The proteomics and metabolomics facilities at UBC are supported by the Canada Foundation for Innovation, the BC Knowledge Development fund, and Genome BC (374PRO). The metabolomics repository is supported by NIH grant U2C-DK119886 and OT2-OD030544.

## References

1. Lopaschuk, G. D. (2017) Metabolic modulators in heart disease: past, present, and future Canadian Journal of Cardiology 33, 838–849

2. De Jong, K. A., and Lopaschuk, G. D. (2017) Complex energy metabolic changes in heart failure with preserved ejection fraction and heart failure with reduced ejection fraction Canadian Journal of Cardiology 33, 860–871

3. Pascual, F., and Coleman, R. A. (2016) Fuel availability and fate in cardiac metabolism: a tale of two substrates Biochimica et Biophysica Acta (BBA)-Molecular and Cell Biology of Lipids 1861, 1425–1433

4. Karwi, Q. G., Uddin, G. M., Ho, K. L., and Lopaschuk, G. D. (2018) Loss of metabolic flexibility in the failing heart Frontiers in cardiovascular medicine 5, 68

5. Randle, P. (1963) The glucose fatty-acid cycle. Its role in insulin sensitivity and the metabolic disturbances of diabetes mellitus Lancet 1, 785–789

6. Peterson, L. R., Herrero, P., Schechtman, K. B., Racette, S. B., Waggoner, A. D., Kisrieva-Ware, Z. et al. (2004) Effect of obesity and insulin resistance on myocardial substrate metabolism and efficiency in young women Circulation 109, 2191–2196

7. Rijzewijk, L. J., van der Meer, R. W., Lamb, H. J., de Jong, H. W., Lubberink, M., Romijn, J. A. et al. (2009) Altered myocardial substrate metabolism and decreased diastolic function in nonischemic human diabetic cardiomyopathy: studies with cardiac positron emission tomography and magnetic resonance imaging Journal of the American College of Cardiology 54, 1524–1532

8. Dávila-Román, V. G., Vedala, G., Herrero, P., De Las Fuentes, L., Rogers, J. G., Kelly, D. P., et al. (2002) Altered myocardial fatty acid and glucose metabolism in idiopathic dilated cardiomyopathy Journal of the American College of Cardiology 40, 271–277

9. Saddik, M., Gamble, J., Witters, L., and Lopaschuk, G. (1993) Acetyl-CoA carboxylase regulation of fatty acid oxidation in the heart Journal of Biological Chemistry 268, 25836–25845

10. Awan, M. M., and Saggerson, E. D. (1993) Malonyl-CoA metabolism in cardiac myocytes and its relevance to the control of fatty acid oxidation Biochemical Journal 295, 61–66

11. Lopaschuk, G. D. (2001) Malonyl CoA control of fatty acid oxidation in the diabetic rat heart In Diabetes and Cardiovascular Disease, Springer, 155–165

12. Randle, P. J. (1998) Regulatory interactions between lipids and carbohydrates: the glucose fatty acid cycle after 35 years Diabetes/metabolism reviews 14, 263–283

13. Sugden, M. C., and Holness, M. J. (2006) Mechanisms underlying regulation of the expression and activities of the mammalian pyruvate dehydrogenase kinases Archives of physiology and biochemistry 112, 139–149

14. Zhang, S., Hulver, M. W., McMillan, R. P., Cline, M. A., and Gilbert, E. R. (2014) The pivotal role of pyruvate dehydrogenase kinases in metabolic flexibility Nutrition & metabolism 11, 1–9

15. Abbot, E. L., McCormack, J. G., Reynet, C., Hassall, D. G., Buchan, K. W., and Yeaman, S. J. (2005) Diverging regulation of pyruvate dehydrogenase kinase isoform gene expression in cultured human muscle cells The FEBS journal 272, 3004–3014

16. Roche, T. a., and Hiromasa, Y. (2007) Pyruvate dehydrogenase kinase regulatory mechanisms and inhibition in treating diabetes, heart ischemia, and cancer Cellular and molecular life sciences 64, 830

17. Popov, K. M., Hawes, J. W., and Harris, R. A. (1997) 9 Mitochondrial α-ketoacid dehydrogenase kinases: A new family of protein kinases Advances in second messenger and phosphoprotein research 31, 105–111

18. Kolobova, E., Tuganova, A., Boulatnikov, I., and Popov, K. M. (2001) Regulation of pyruvate dehydrogenase activity through phosphorylation at multiple sites Biochemical Journal 358, 69–77

19. Korotchkina, L. G., and Patel, M. S. (2001) Site specificity of four pyruvate dehydrogenase kinase isoenzymes toward the three phosphorylation sites of human pyruvate dehydrogenase Journal of Biological Chemistry 276, 37223–37229

20. Bowker-Kinley, M. M., Davis, I. W., Wu, P., Harris, A. R., and Popov, M. K. (1998) Evidence for existence of tissue-specific regulation of the mammalian pyruvate dehydrogenase complex Biochemical Journal 329, 191–196

21. Harris, R. A., Huang, B., and Wu, P. (2001) Control of pyruvate dehydrogenase kinase gene expression Advances in enzyme regulation 41, 269–288

22. Peters, S. (2003) Regulation of PDH activity and isoform expression: diet and exercise Portland Press Ltd.,

23. Sugden, M. C., Bulmer, K., Augustine, D., and Holness, M. J. (2001) Selective modification of pyruvate dehydrogenase kinase isoform expression in rat pancreatic islets elicited by starvation and activation of peroxisome proliferator–activated receptor-α: implications for glucose-stimulated insulin secretion Diabetes 50, 2729–2736

24. Members, C.-N., and Partners (2024) Database Resources of the National Genomics Data Center, China National Center for Bioinformation in 2025 Nucleic Acids Research 10.1093/nar/gkae978

25. Chen, L., Yang, F., Chen, X., Rao, M., Zhang, N.-N., Chen, K. et al. (2017) Comprehensive myocardial proteogenomics profiling reveals C/EBPα as the key factor in the lipid storage of ARVC Journal of Proteome Research 16, 2863–2876

26. Liu, S., Xia, Y., Liu, X., Wang, Y., Chen, Z., Xie, J., et al. (2017) In-depth proteomic profiling of left ventricular tissues in human end-stage dilated cardiomyopathy Oncotarget 8, 48321

27. Yi, X., Jiang, D.-S., Feng, G., Jiang, X.-j., and Zeng, H.-L. (2019) An altered left ventricle protein profile in human ischemic cardiomyopathy revealed in comparative quantitative proteomics Kardiologia Polska (Polish Heart Journal) 77, 951–959

28. Tromp, J., Westenbrink, B. D., Ouwerkerk, W., van Veldhuisen, D. J., Samani, N. J., Ponikowski, P. et al. (2018) Identifying pathophysiological mechanisms in heart failure with reduced versus preserved ejection fraction Journal of the American College of Cardiology 72, 1081–1090

29. Reitz, C. J., Tavassoli, M., Kim, D. H., Shah, S., Lakin, R., Teng, A. C. et al. (2023) Proteomics and phosphoproteomics of failing human left ventricle identifies dilated cardiomyopathy-associated phosphorylation of CTNNA3 Proceedings of the National Academy of Sciences 120, e2212118120

30. Stoehr, J. P., Byers, J. E., Clee, S. M., Lan, H., Boronenkov, I. V., Schueler, K. L. et al. (2004) Identification of major quantitative trait loci controlling body weight variation in ob/ob mice Diabetes 53, 245–249

31. Leung, C. L., Karunakaran, S., Atser, M. G., Innala, L., Hu, X., Viau, V. et al. (2023) Analysis of a genetic region affecting mouse body weight Physiological genomics 55, 132–146

32. Branco, A. F., Pereira, S. P., Gonzalez, S., Gusev, O., Rizvanov, A. A., and Oliveira, P. J. (2015) Gene expression profiling of H9c2 myoblast differentiation towards a cardiac-like phenotype PloS one 10, e0129303

33. Fukushima, A., Zhang, L., Huqi, A., Lam, V. H., Rawat, S., Altamimi, T., et al. (2018) Acetylation contributes to hypertrophy-caused maturational delay of cardiac energy metabolism JCI insight 3,

34. Opie, L. H. (2014) Cardiac metabolism in health and disease In Cellular and Molecular Pathobiology of Cardiovascular Disease, Elsevier, 23–36

35. Wiese, E. K., Hitosugi, S., Loa, S. T., Sreedhar, A., Andres-Beck, L. G., Kurmi, K. et al. (2021) Enzymatic activation of pyruvate kinase increases cytosolic oxaloacetate to inhibit the Warburg effect Nature metabolism 3, 954–968

36. Gray, L. R., Tompkins, S. C., and Taylor, E. B. (2014) Regulation of pyruvate metabolism and human disease Cellular and molecular life sciences 71, 2577–2604

37. Sugden, M. C., Kraus, A., Harris, R. A., and Holness, M. J. (2000) Fibre-type specific modification of the activity and regulation of skeletal muscle pyruvate dehydrogenase kinase (PDK) by prolonged starvation and refeeding is associated with targeted regulation of PDK isoenzyme 4 expression Biochemical Journal 346, 651–657

38. Iso, T., and Kurabayashi, M. (2017) Fatty Acid Uptake by the Heart During Fasting In Handbook of Famine, Starvation, and Nutrient Deprivation: From Biology to Policy, Preedy V, and Patel VB, eds. Springer International Publishing, Cham 1–20

39. Wunderling, K., Zurkovic, J., Zink, F., Kuerschner, L., and Thiele, C. (2023) Triglyceride cycling enables modification of stored fatty acids Nature Metabolism 5, 699–709

40. Alabed, H. B., Mancini, D. F., Buratta, S., Calzoni, E., Giacomo, D. D., Emiliani, C. et al. (2024) LipidOne 2.0: a web tool for discovering biological meanings hidden in lipidomic data Current Protocols 4, e70009

41. Priestman, D. A., Orfali, K. A., and Sugden, M. C. (1996) Pyruvate inhibition of pyruvate dehydrogenase kinase: effects of progressive starvation and hyperthyroidism in vivo, and of dibutyryl cyclic AMP and fatty acids in cultured cardiac myocytes FEBS letters 393, 174–178

42. Marchington, D. R., Kerbey, A. L., and Randle, P. J. (1990) Longer-term regulation of pyruvate dehydrogenase kinase in cultured rat cardiac myocytes Biochemical Journal 267, 245–247

43. Khan, A. U. H., Salehi, H., Alexia, C., Valdivielso, J. M., Bozic, M., Lopez-Mejia, I. C. et al. (2022) Glucose starvation or pyruvate dehydrogenase activation induce a broad, ERK5-mediated, metabolic remodeling leading to fatty acid oxidation Cells 11, 1392

44. Cardoso, A. C., Lam, N. T., Savla, J. J., Nakada, Y., Pereira, A. H. M., Elnwasany, A. et al. (2020) Mitochondrial substrate utilization regulates cardiomyocyte cell-cycle progression Nature metabolism 2, 167–178

45. Klyuyeva, A., Tuganova, A., Kedishvili, N., and Popov, K. M. (2019) Tissue-specific kinase expression and activity regulate flux through the pyruvate dehydrogenase complex. Journal of Biological Chemistry 294, 838–851

46. Semba, H., Takeda, N., Isagawa, T., Sugiura, Y., Honda, K., Wake, M., et al. (2016) HIF-1α-PDK1 axis-induced active glycolysis plays an essential role in macrophage migratory capacity Nature communications 7, 11635

47. Li, X., Jiang, Y., Meisenhelder, J., Yang, W., Hawke, D. H., Zheng, Y. et al. (2016) Mitochondria-translocated PGK1 functions as a protein kinase to coordinate glycolysis and the TCA cycle in tumorigenesis Molecular cell 61, 705–719

48. Oh, S., Mai, X. L., Kim, J., de Guzman, A. C. V., Lee, J. Y., and Park, S. (2024) Glycerol 3-phosphate dehydrogenases (1 and 2) in cancer and other diseases Experimental & Molecular Medicine 1–14

49. Zechner, R., Madeo, F., and Kratky, D. (2017) Cytosolic lipolysis and lipophagy: two sides of the same coin Nature Reviews Molecular Cell Biology 18, 671–684

50. Banke, N. H., Wende, A. R., Leone, T. C., O’Donnell, J. M., Abel, E. D., Kelly, D. P. et al. (2010) Preferential oxidation of triacylglyceride-derived fatty acids in heart is augmented by the nuclear receptor PPARα Circulation research 107, 233–241

51. Kienesberger, P. C., Pulinilkunnil, T., Nagendran, J., and Dyck, J. R. (2013) Myocardial triacylglycerol metabolism Journal of molecular and cellular cardiology 55, 101–110

52. Swanton, M. E., and Saggerson, E. D. (1997) Effects of adrenaline on triacylglycerol synthesis and turnover in ventricular myocytes from adult rats Biochemical Journal 328, 913–922

53. Zimmermann, R., Strauss, J. G., Haemmerle, G., Schoiswohl, G., Birner-Gruenberger, R., Riederer, M. et al. (2004) Fat mobilization in adipose tissue is promoted by adipose triglyceride lipase Science 306, 1383–1386

54. Kloska, A., Węsierska, M., Malinowska, M., Gabig-Cimińska, M., and Jakóbkiewicz-Banecka, J. (2020) Lipophagy and lipolysis status in lipid storage and lipid metabolism diseases International journal of molecular sciences 21, 6113

55. Singh, R., Kaushik, S., Wang, Y., Xiang, Y., Novak, I., Komatsu, M. et al. (2009) Autophagy regulates lipid metabolism Nature 458, 1131–1135

56. Rambold, A. S., Cohen, S., and Lippincott-Schwartz, J. (2015) Fatty acid trafficking in starved cells: regulation by lipid droplet lipolysis, autophagy, and mitochondrial fusion dynamics Developmental cell 32, 678–692

57. Jovičić, E. J., Janež, A. P., Eichmann, T. O., Koren, Š., Brglez, V., Jordan, P. M., et al. (2023) Lipid droplets control mitogenic lipid mediator production in human cancer cells Molecular metabolism 76, 101791

58. Sathyanarayan, A., Mashek, M. T., and Mashek, D. G. (2017) ATGL promotes autophagy/lipophagy via SIRT1 to control hepatic lipid droplet catabolism Cell reports 19, 1–9

59. Najt, C. P., Devarajan, M., and Mashek, D. G. (2022) Perilipins at a glance Journal of cell science 135, jcs259501

60. Keenan, S. N., De Nardo, W., Lou, J., Schittenhelm, R. B., Montgomery, M. K., Granneman, J. G. et al. (2021) Perilipin 5 S155 phosphorylation by PKA is required for the control of hepatic lipid metabolism and glycemic control Journal of lipid research 62,

61. Wang, H., Sreenivasan, U., Hu, H., Saladino, A., Polster, B. M., Lund, L. M. et al. (2011) Perilipin 5, a lipid droplet-associated protein, provides physical and metabolic linkage to mitochondria Journal of lipid research 52, 2159–2168

62. Thiele, C., Papan, C., Hoelper, D., Kusserow, K., Gaebler, A., Schoene, M. et al. (2012) Tracing fatty acid metabolism by click chemistry ACS chemical biology 7, 2004–2011

63. Demichev, V., Yu, F., Teo, G. C., Szyrwiel, L., Rosenberger, G. A., Decker, J. et al. (2021) High sensitivity dia-PASEF proteomics with DIA-NN and FragPipe BioRxiv

64. Cajka, T., Smilowitz, J. T., and Fiehn, O. (2017) Validating quantitative untargeted lipidomics across nine liquid chromatography–high-resolution mass spectrometry platforms Analytical chemistry 89, 12360–12368

65. Qu, X., Bhalla, K., Horianopoulos, L. C., Hu, G., Alcázar Magaña, A., Foster, L. J., et al. (2024) Phosphate availability conditions caspofungin tolerance, capsule attachment and titan cell formation in Cryptococcus neoformans Frontiers in Fungal Biology 5, 1447588

66. Tsugawa, H., Cajka, T., Kind, T., Ma, Y., Higgins, B., Ikeda, K. et al. (2015) MS-DIAL: data-independent MS/MS deconvolution for comprehensive metabolome analysis Nature Methods 12, 523–526

67. Black, B., da Silva, L. B. R., Hu, G., Qu, X., Smith, D. F., Magaña, A. A., et al. (2024) Glutathione-mediated redox regulation in Cryptococcus neoformans impacts virulence Nature microbiology 1–15

68. McAfee, A., Magana, A. A., Foster, L. J., and Hoover, S. E. (2024) Parallel pheromone, metabolite, and lipid analyses reveal patterns associated with early life transitions and ovary activation in honey bee (Apis mellifera) queens bioRxiv 2024.2004. 2019.590367

69. Adams, K. J., Pratt, B., Bose, N., Dubois, L. G., St. John-Williams, L., Perrott, K. M. et al. (2020) Skyline for small molecules: a unifying software package for quantitative metabolomics Journal of proteome research 19, 1447–1458

70. Heinrich, P., Kohler, C., Ellmann, L., Kuerner, P., Spang, R., Oefner, P. J. et al. (2018) Correcting for natural isotope abundance and tracer impurity in MS-, MS/MS-and high-resolution-multiple-tracer-data from stable isotope labeling experiments with IsoCorrectoR Scientific reports 8, 17910

71. Ritchie, M. E., Phipson, B., Wu, D., Hu, Y., Law, C. W., Shi, W., et al. (2015) limma powers differential expression analyses for RNA-sequencing and microarray studies Nucleic acids research 43, e47–e47

72. Sud, M., Fahy, E., Cotter, D., Azam, K., Vadivelu, I., Burant, C. et al. (2016) Metabolomics Workbench: An international repository for metabolomics data and metadata, metabolite standards, protocols, tutorials and training, and analysis tools Nucleic acids research 44, D463–D470

