## Supplemental figures for "Pyruvate dehydrogenase kinase 1 controls triacylglycerol hydrolysis in cardiomyocytes"

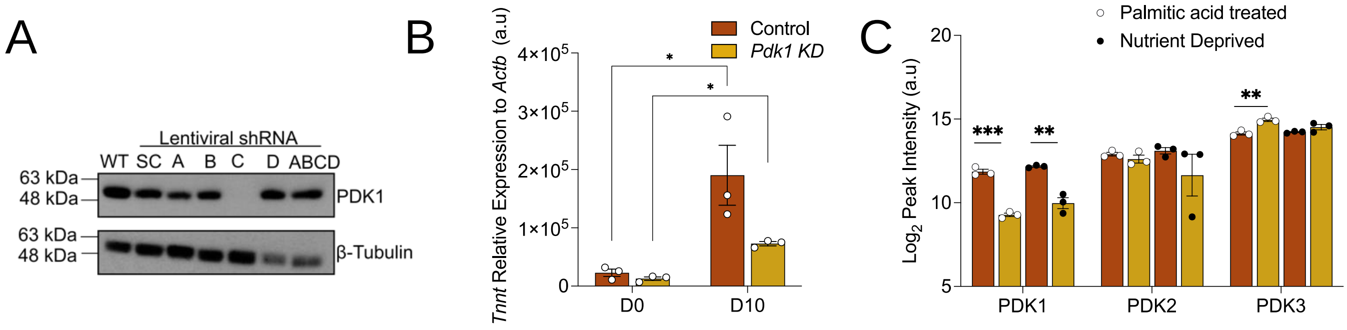


**Figure S1: Model Validation. (A)** Representative Western blot of PDK1 protein abundance following transduction of H9c2 myoblast with four lentiviral shRNA (A, B, C (*Pdk1* KD), D) targeting *Pdk1* and a scrambled shRNA (SC) as control. Untransduced cells are denoted as wildtype (WT). **(B)** Cardiac troponin T (*Tnnt*) gene expression relative to β-actin (*Actb*) in differentiated H9c2 cardiomyocytes measured by reverse-transcription PCR. **(C)** PDK protein abundance measured by LC-MS/MS based proteomics in control and *Pdk1* knockdown (KD) differentiated H9c2 cardiomyocytes following 375 μM palmitic acid treatment and nutrient deprivation in 5.5 mM glucose, serum-free culture media. Data are shown as mean ± SEM and were analyzed by a two-tailed t-test. * $p < 0.05$; ** $p < 0.01$; *** $p < 0.001$; **** $p < 0.0001$.

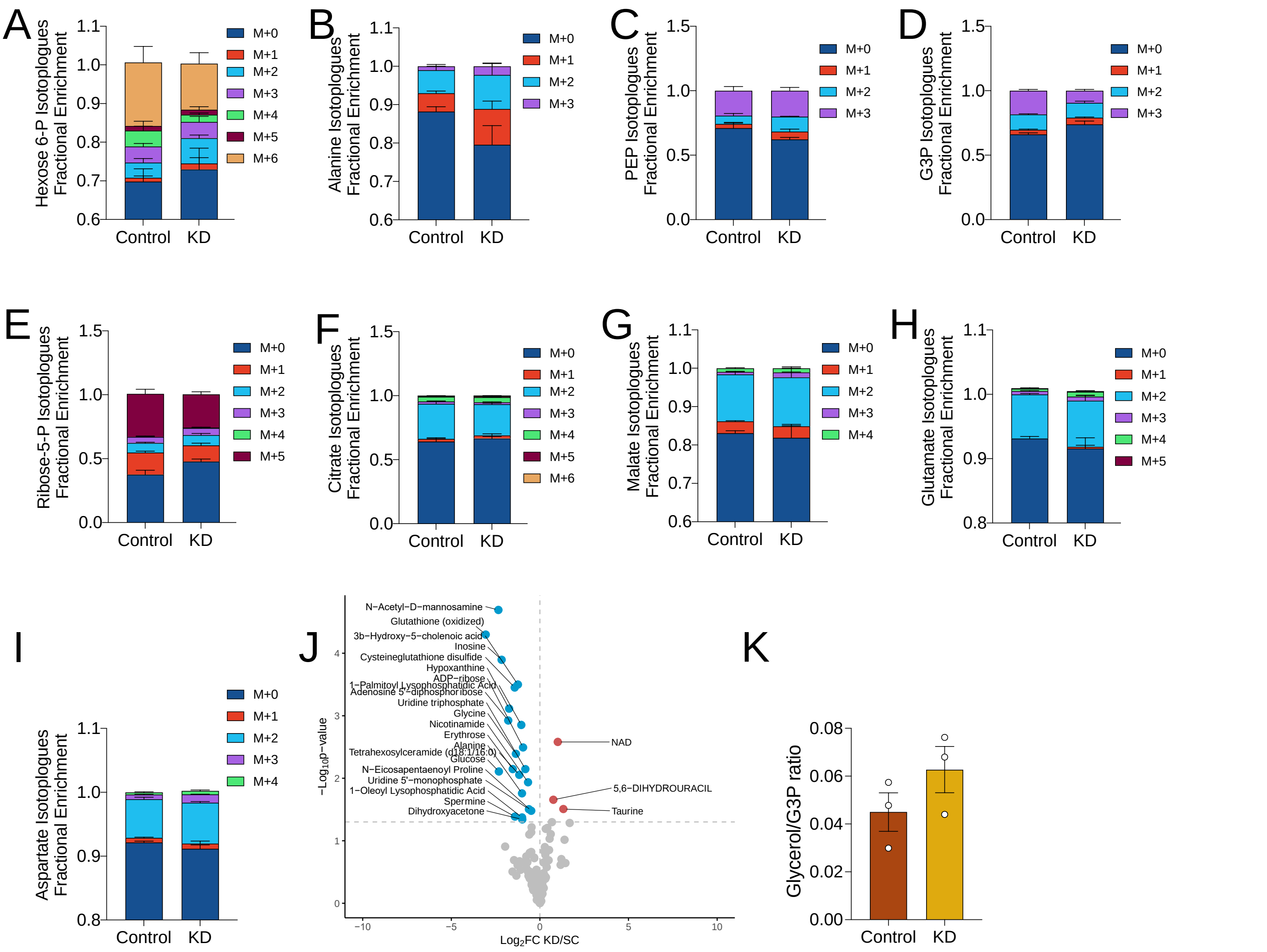


**Figure S2: Metabolomics.** Cells were treated with palmitic acid and nutrient deprived in low glucose, serum-free media containing 5.5 mM of $[U\text{-}^{13}C_6]$ glucose for 1 h. Fractional enrichments of **(A)** hexose-6-phosphate, **(B)** alanine, **(C)** phosphoenolpyruvate, **(D)** glyecerol-3-phosphate, **(E)** ribose-5-phosphate, **(F)** citrate, **(G)** malate, **(H)** glutamate, and **(I)** aspartate isotopologues were then assessed. **(J)** Unlabelled glycerol to glycerol-3-phosphate ratio was assessed in unlabelled cells. **(K)** Volcano plot of identified metabolites in cells following 3 h nutrient deprivation. Blue and red dots signify downregulated and upregulated metabolites in *Pdk1* knockdown cells relative to control. Data are shown as mean ± SEM and were analyzed by Limma.

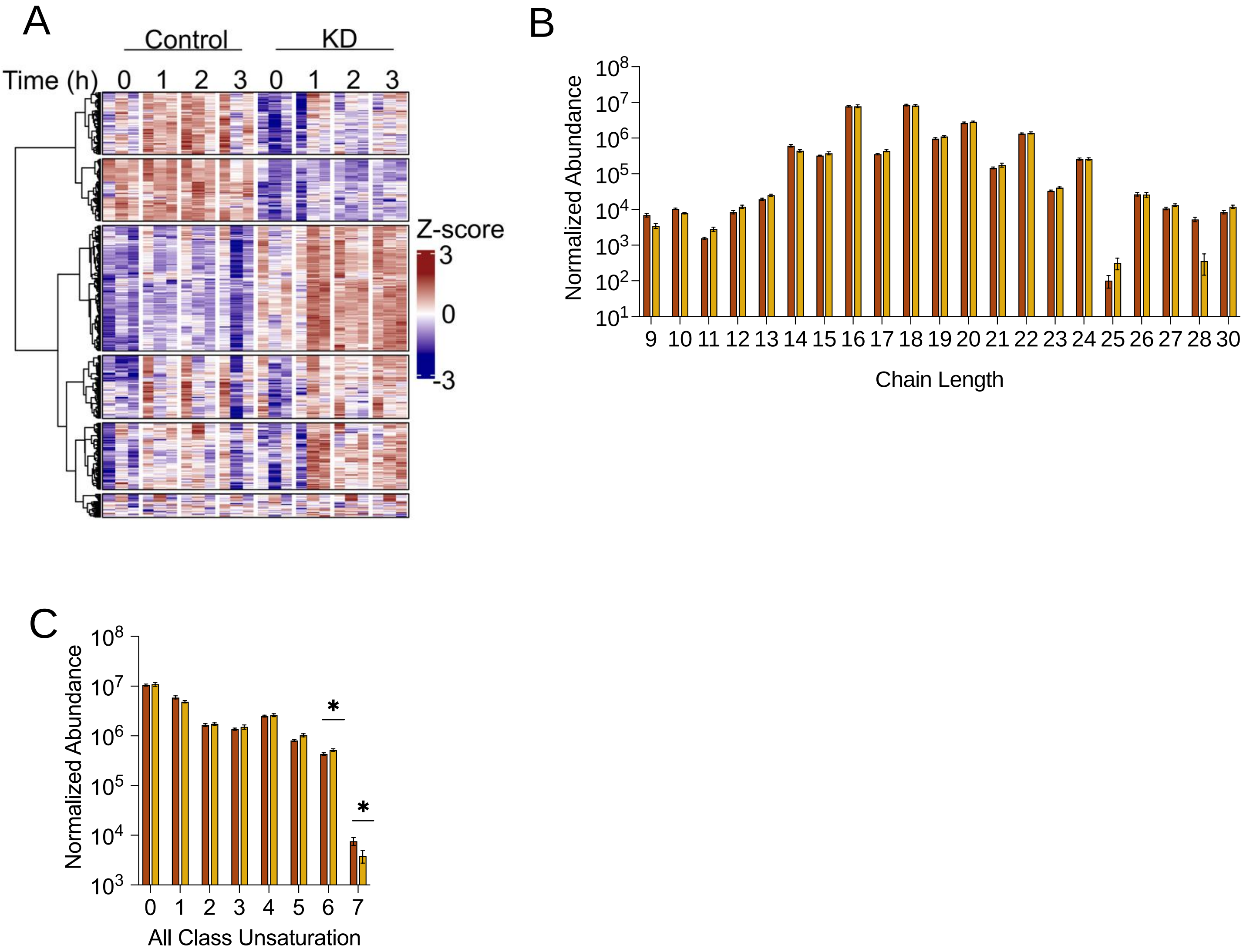


**Figure S3: Lipidomics. (A)** Heatmap of identified lipids from cells treated with 150 μM palmitic acid and nutrient deprived in low glucose, serum-free media for 3 h. Sum abundance of all lipid species of certain **(B)** chain length and **(C)** unsaturation in cells following palmitic acid treatment**.** Data are shown as mean ± SEM and were analyzed by multiple t-test with Bonferroni-Dunn correction. * $p < 0.05$; ** $p < 0.01$; *** $p < 0.001$; **** $p < 0.0001$.

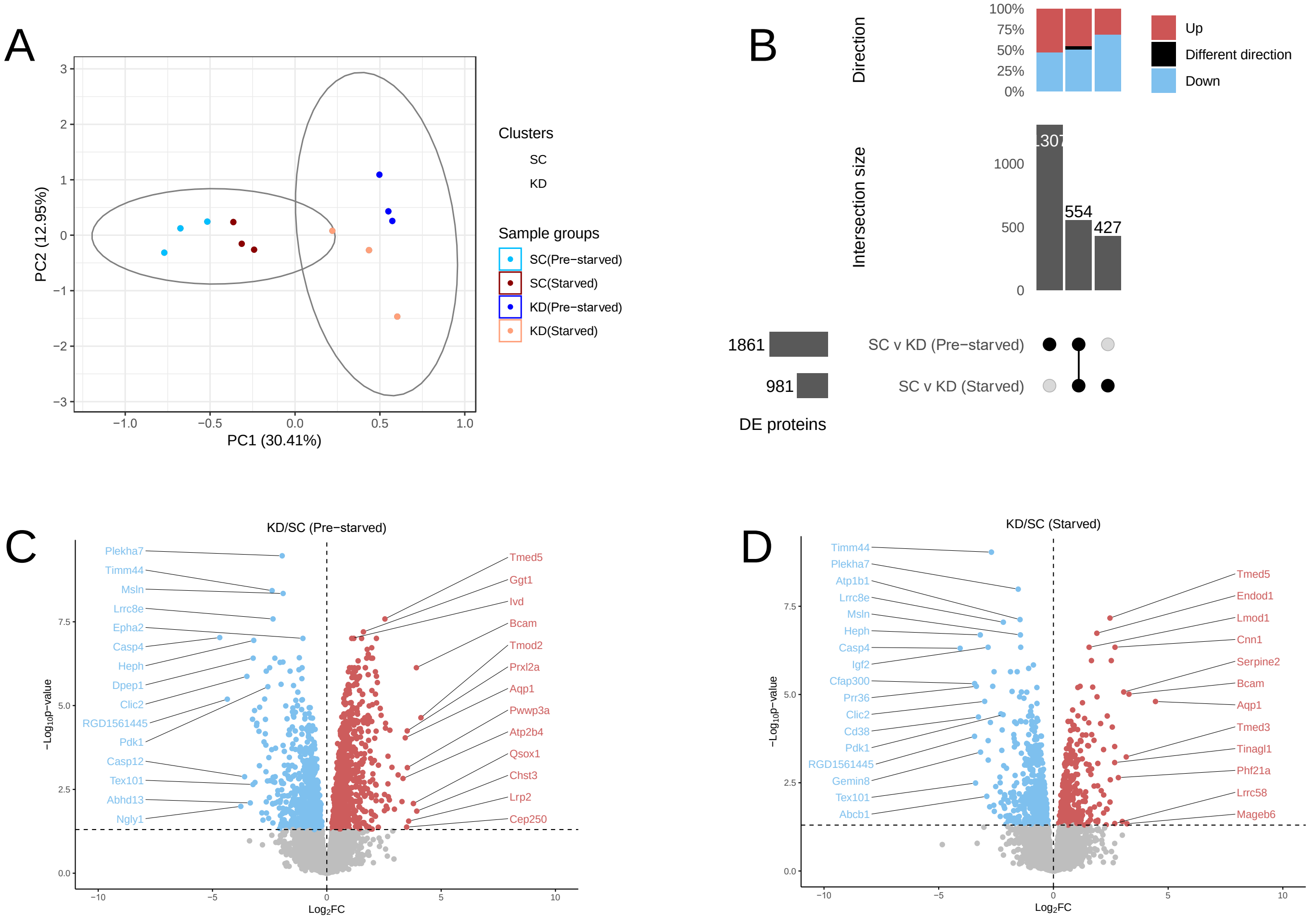


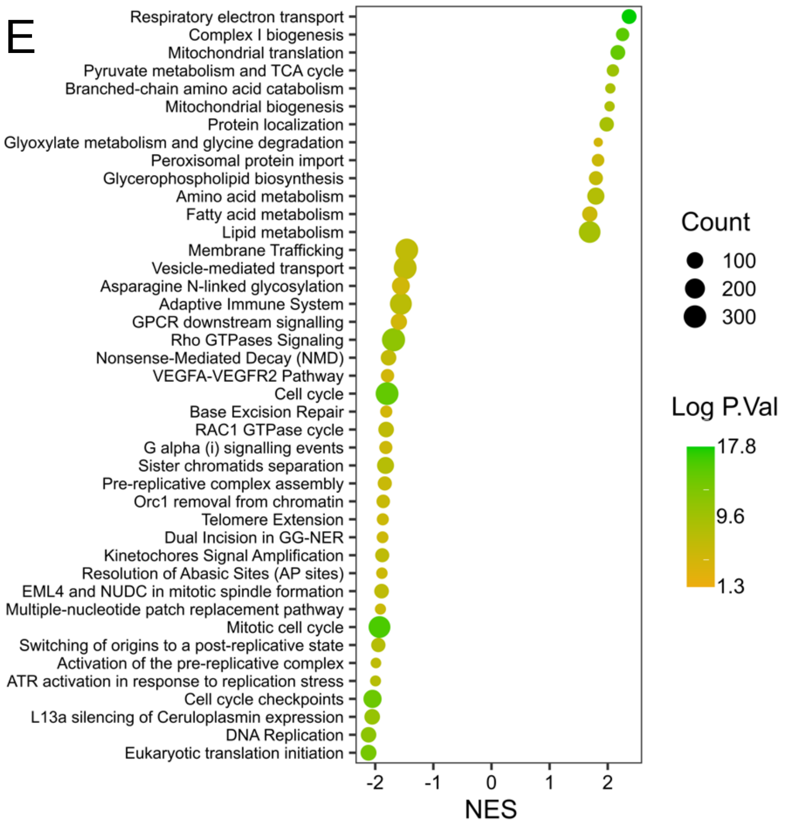


**Figure S4: Proteomics.** Mass-spec based proteomics was run on cells treated with 375 μM palmitic acid for 18 h (pre-starved), then nutrient deprived in low glucose, serum-free media (starved) for 3 h. Identified proteins were analyzed by **(A)** Principal component analysis, **(B)** UpSet plot to show the intersection of differentially expressed proteins in *Pdk1* knockdown cells relative to control in pre-starved and starved conditions, and volcano plot in **(C)** pre-starved and **(D)** starved conditions. **(E)** Pathway enrichment analysis was then carried out on differentially expressed proteins in *Pdk1* knockdown cells relative to control based on gene-set enrichment analysis derived from the Reactome database.

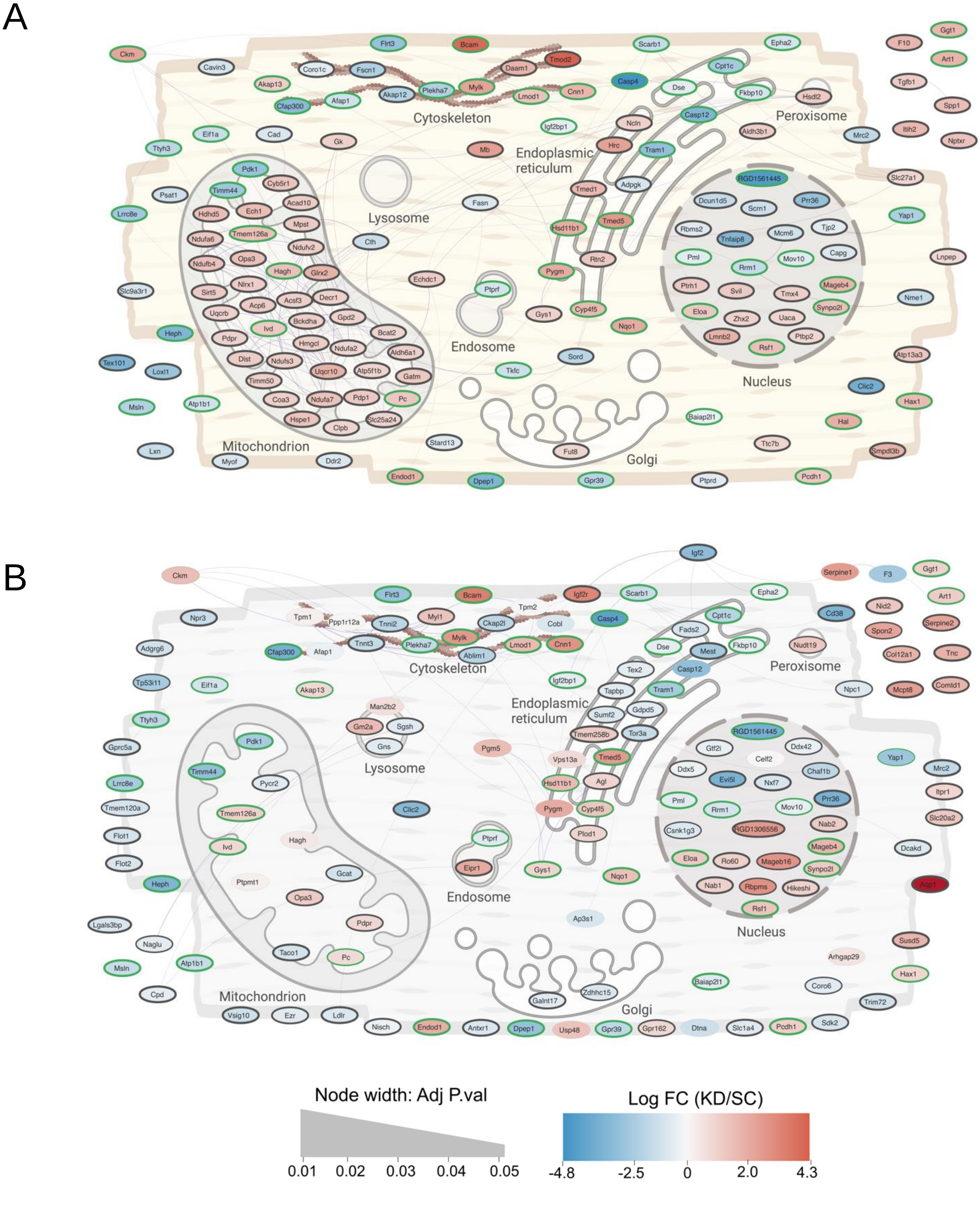


**Figure S5: Subcellular mapping of top 150 proteins.** Highest confidence (top 150 lowest p-values, per condition) differentially abundant proteins between control and *Pdk1* knockdown cells treated with palmitic acid were connected in a medium-confidence (functional score 0.50) protein-protein interaction network (purple lines) using STRING and depicted in the context of their sub-cellular locations. The color of the nodes represents the fold change in *Pdk1* knockdown cells relative to control, while the thickness of the line around the nodes represents the adjusted p-value, and a green node border illustrates a common protein present in both conditions.

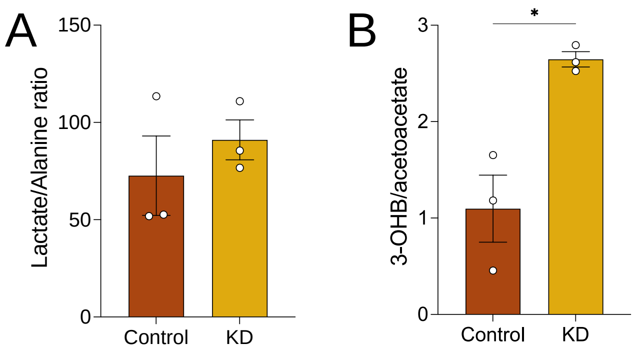


**Figure S6: Assessments of Cytosolic and Mitochondrial Redox States.** Cells were treated with palmitic acid and nutrient deprived in low glucose for 3 h. Metabolite ratios using low confidence-scored annotated metabolites. A high confidence threshold was set at a Progeneis QI score of ≥45. (A) Lactate (score = 40.3) to alanine (score = 49.3) ratio to assess cytosolic redox state and (B) 3-hydroxybutyrate (score = 40) to acetoacetate (score = 36.3) to assess mitochondrial redox state. Data are shown as mean ± SEM and were analyzed by Two-tailed t-test. * $p < 0.05$; ** $p < 0.01$; *** $p < 0.001$; **** $p < 0.0001$.
